# Comprehensive degron mapping in human transcription factors

**DOI:** 10.1101/2025.05.16.654404

**Authors:** Fia B. Larsen, Vasileios Voutsinos, Nicolas Jonsson, Kristoffer E. Johansson, Freia D. Ethelberg, Kresten Lindorff-Larsen, Rasmus Hartmann-Petersen

**Author notes:** Corresponding authors: V.V., K.L.-L., R.H.-P.

## Abstract

Gene expression is regulated by the targeted degradation of transcription factors through the ubiquitin-proteasome system. Transcription factors destined for degradation are recognized by E3 ubiquitin-protein ligases through short motifs termed degrons, embedded within the sequence. In this study, we systematically map degrons in all 1,626 human transcription factors. We find thousands of both known and previously unidentified degrons and characterize their sequence properties. Degrons placed within exposed and intrinsically disordered regions regulate the cellular abundance of the transcription factors, while the most common somatic mutations that are linked to skin cutaneous melanoma lead to unfolding and exposure of a buried degron in zinc fingers. We present examples of compartment specific degrons and demonstrate that variant effects in transcription factors correlate with degron potency. Finally, we show that while >60% of all predicted transcriptional activation domains overlap with strong degrons, acidic residues within the remaining transactivating regions counter the degron potency.

## Introduction

The proteome exists in a finely balanced state of homeostasis (proteostasis), where proteins are constantly degraded and replaced by newly synthesized ones (1, 2). Intracellular protein degradation is primarily mediated by the ubiquitin-proteasome system (UPS), which also indirectly regulates protein synthesis through targeted degradation of transcription factors (TFs) (3-5). Many TFs have been found to be short-lived, which allows for a tight regulation of gene expression, critical for a wide range of cellular processes (4, 6).

Although a few proteins, including some TFs, have been reported to be degraded by the proteasome independently of ubiquitin (7, 8), most proteins, including many TFs, are UPS targets (3-5). The specificity of the UPS is in general governed by ∼600 different E3 ubiquitin-protein ligases that catalyze ubiquitylation of target proteins (9-11). Once ubiquitylated, the proteins gain affinity for the 26S proteasome, which then threads the target proteins into the proteasome lumen, where the proteolysis occurs (12). The features that are recognized by the E3s are the so-called degradation signals or “degrons” (13-15), which are typically short motifs embedded within the amino acid sequence of the target protein. Well-characterized examples include the N- and C-degrons (16, 17), where specific amino acid residues near the N- or C-terminus of the target proteins, are recognized by certain E3s. Some degrons, such as the KEN box or destruction box of cyclins (18), the AUX degron (19), and phospho-degrons (10) are regulated by activation of the E3s, small molecules or post-translational modifications. Finally, protein quality control (PQC) degrons, which typically comprise hydrophobic sequences buried within the core of a folded protein (20-23), only become active upon exposure such as during an unfolding or misfolding event (24), thus ensuring the selective degradation of non-native proteins.

The proteasome and most other UPS components are found in both the cytosol and nucleus (25-27). However, as certain E3s have been shown to localize exclusively to one of these compartments, some differences in degron recognition and degradation are expected between these compartments. For instance, since the E3 complex CRL4 with its substrate receptor DCAF12 is known to localize to the nucleus (28), possibly DCAF12 degrons are only relevant in nuclear target proteins. Similarly, proteins that require binding to nuclear PCNA for degradation (29) are also expected to display nuclear specific degradation. Conversely, PQC degrons seem to be broadly active (30, 31), albeit studies in yeast have shown that certain misfolded proteins are translocated to the nucleus prior to their degradation (32, 33). Accordingly, a major challenge is to determine to what extent different compartments impose constraints on degron activity.

To further our general understanding of proteostasis and to gain a systems-level understanding of the UPS, it is essential to map degrons in a wide variety of relevant target proteins. Considering the size of the proteome, this is a painstaking endeavor, which has only recently become possible through the development of high throughput screening methods. Degron mapping by such technologies (20-22, 30, 34-37), has resulted in important insights into PQC (20, 21, 30) and cullin-RING ligase (CRL) degrons (20, 21), and allowed development of computational tools for degron prediction (21, 37, 38). It has also led to the discovery of novel degradation pathways (8, 20, 36, 39). However, much remains to be learned, particularly concerning compartment-specific degradation, as well as degron formation or disruption by mutations. Also the relevance of degron positioning and exposure in structured versus intrinsically disordered regions (IDRs) needs to be explored in more detail.

Here, we focus on transcription factors. Combining tiled peptide libraries with flow sorting and sequencing, we systematically mapped thousands of degrons in 1,626 different human TFs. For selected degrons in exposed IDRs, we show their relevance for regulating the abundance of the full-length proteins. We show that the most common somatic mutations linked to skin cutaneous melanoma (40), lead to exposure of PQC degrons locally embedded in C2H2 Zn finger proteins. Finally, we present examples of compartment-specific degrons and show that many transactivating regions also function as degrons. The dataset provides a rich resource for future in-depth studies of individual TFs and underlines the importance of proteostasis for transcriptional control.

## Results

### High throughput screening for degrons in human transcription factors

To map degrons in human TFs, 1,626 different human TFs (41) were divided into 30-residue tiles, overlapping by 15 residues, resulting in a tiled TF library encompassing 62,992 tiles (**Fig. 1A**). We hypothesized that 30 residues would be sufficiently long to contain a degron, but in most cases too short to form stable protein structures. To avoid potential problems with template switching during PCR and to reduce the complexity of the assayed library, the library was divided into three sub-groups, termed the odds (31,080 tiles), evens (30,292 tiles) and C-terminal (1,620 tiles) libraries. The libraries were then cloned into an integrative mammalian expression vector (42) fused to the C-terminus of GFP (**Fig. 1A**). To focus on TF degradation in the nucleus, the GFP carried an N-terminal nuclear localization signal (NLS). To account for cell-to-cell variations, a reporter protein, mCherry, was co-translationally expressed from an internal ribosome entry site (IRES) (**Fig. 1A**). The plasmid was introduced into HEK293T cells containing a Bxb1 landing pad site (43), thus allowing site-specific integration of the plasmid. As the plasmid lacks a promoter, any non-integrated plasmids should not be expressed. Since correct integration of the plasmid in the landing pad will displace the BFP-iCasp9-Blast^R^ cassette (**Fig. 1A**), non-recombinant cells can be eliminated from the culture using AP1903, which induces apoptosis of iCasp9 positive cells. Thus, similar to other experimental systems (20, 30), GFP fluorescence, normalized to the mCherry signal, will report on the abundance of the GFP-tile fusion and thus reflect the tile’s degron potency.

**Figure 1.**
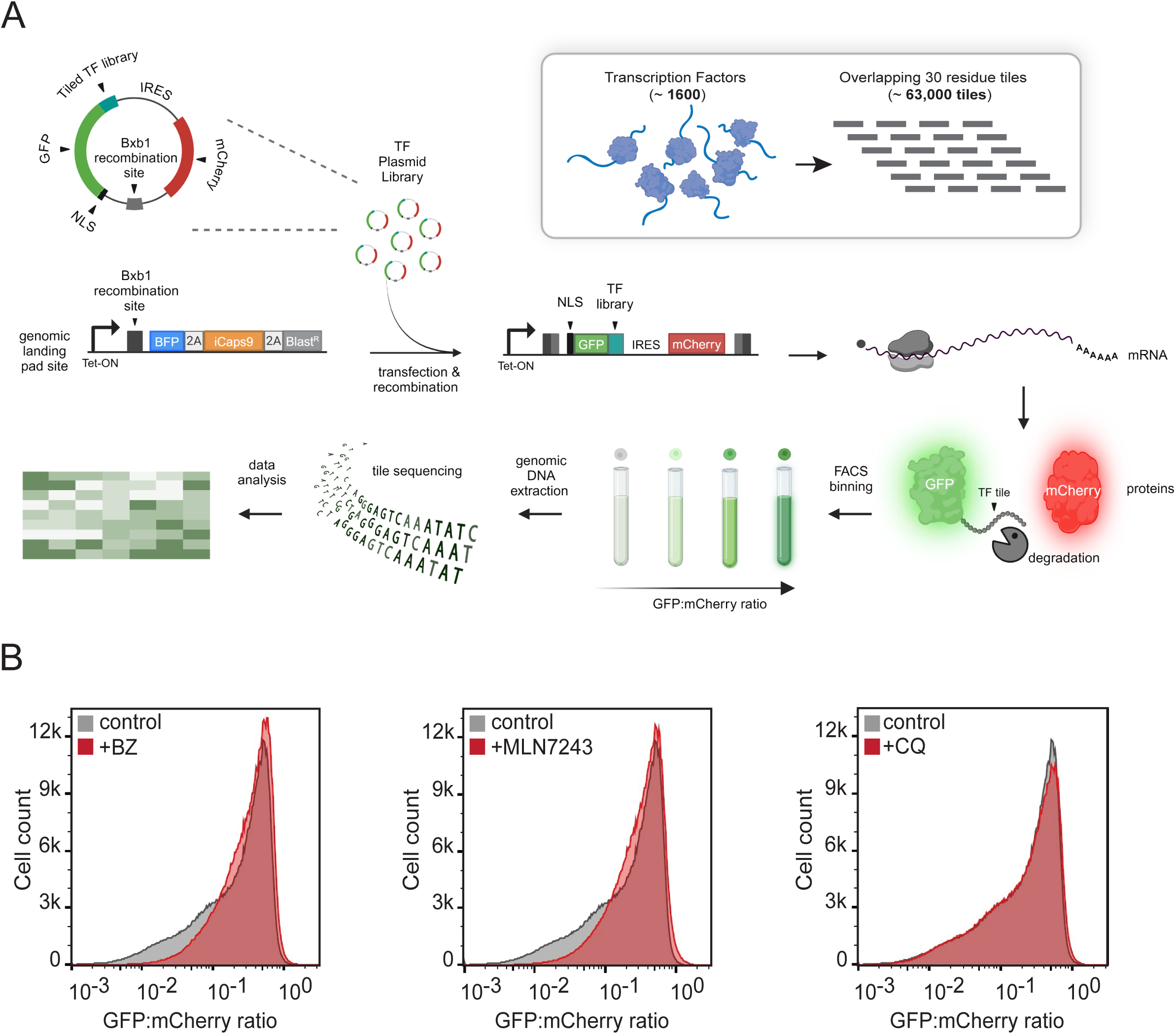
Tiled transcription factor library and degron screening. (A) Schematic representation of the utilized expression system and degron screen. A library of all human transcription factors (TFs) (1626 proteins) was divided into 30-residue tiles overlapping with 15 residues (62,992 tiles). The tiled library was cloned into an integrative mammalian expression vector fused to GFP, containing an N-terminal nuclear localization signal (NLS), followed by the reporter protein mCherry downstream of an internal ribosome entry site (IRES). Upon successful Bxb1-mediated site-specific integration of the plasmid DNA into a landing pad in the HEK293T genome, the GFP-TF library-fusions and mCherry are synthesized from the same mRNA. Non-recombinant cells will express BFP, iCasp9 and blasticidin resistance (Blast^R^) genes, separated by two 2A-like translational start-stop sequences (2A). Cells were sorted into four bins based on the GFP:mCherry ratios using fluorescence activated cell sorting (FACS). Subsequently, the genomic DNA was extracted and sequenced. The figure was created with BioRender.com adapted from (62, 63). (B) Representative flow cytometry profiles displaying the quantification of the GFP:mCherry ratio of cells expressing the TF odd sublibrary untreated control (grey) (n=6×10^5^) or treated for 16 hours with 15 μM bortezomib (BZ) (n=6×10^5^) (red), 1 μM MLN7243 (n=6×10^5^) (red) or 20 µM chloroquine (CQ) (n=6×10^5^) (red).

Initial analysis of the three sublibraries by flow cytometry revealed a similar distribution in GFP:mCherry ratios (**Fig. 1B, Fig. S1**). The majority of the population displayed high GFP:mCherry ratios, corresponding to tiles with no or only minor degron activity. However, a population with lower GFP:mCherry ratios was visible as a shoulder leaning towards reduced GFP:mCherry ratios (**Fig. 1B**), likely corresponding to tiles with increasing degron activity. To investigate if the low abundance was due to ubiquitin-dependent proteasomal degradation, cells were treated with proteasome inhibitor bortezomib (BZ) or an inhibitor of the E1 ubiquitin-activating enzyme (MLN7243). Both treatments resulted in a pronounced shift of the low abundance tiles towards increased GFP:mCherry ratios (**Fig. 1B**). In addition, treatment with cycloheximide (CHX), which inhibits translation, resulted in an increased population of cells displaying low GFP:mCherry ratios (**Fig. S2A**). Conversely, treatment with the autophagy inhibitor chloroquine (CQ), did not affect the GFP:mCherry ratios of the tiles (**Fig. 1B**). This indicates that most of the tiles displaying a reduced abundance contain degrons for ubiquitin-dependent proteasomal degradation, while autophagy does not strongly contribute to the degradation. Since the family of CRLs constitutes a large and important group of E3s, we also analyzed the distribution of the library in cultures with the NEDD8 E1 inhibitor (MLN4924). Since this resulted in a modest increase in abundance (**Fig. S2B**), it is likely that only a subset of the degrons depend on CRLs.

### Library screening and validation

To identify degrons, we used fluorescence activated cell sorting (FACS) to separate the cells based on the GFP:mCherry ratios into four bins, each containing 25% of the population (**Fig. 2A**). Then, the frequency of the individual tiles in each bin was determined by extracting genomic DNA from the sorted cells, followed by sequencing across the 30 residue (90 bp) tiles. In turn, this allowed us to assign each of the tiles with a degron score (see Methods). Degron scores close to zero indicate low degron potency, while scores close to one indicate high degron potency. The calculated degron scores are based on three biological replicates (separate transfections), each with two FACS replicates. The degron scores correlated well between replicates (average Pearson correlation of 0.97) (**Fig. S3-S5**). We attained data for 58,781 tiles out of the initial 63,011 tiles (incl. control peptides), corresponding to a total coverage of 93% of the entire TF proteome. Due to the utilized gating scheme, the degron scores are distributed in four peaks (**Fig. 2B**) with 45% of the library having a degron score >0.5. The degron scores obtained from the screen correlated strongly with low throughput flow cytometry analysis of cells transfected individually with 82 different tiles (Pearson correlation -0.93) (**Fig. 2B**).

**Figure 2.**
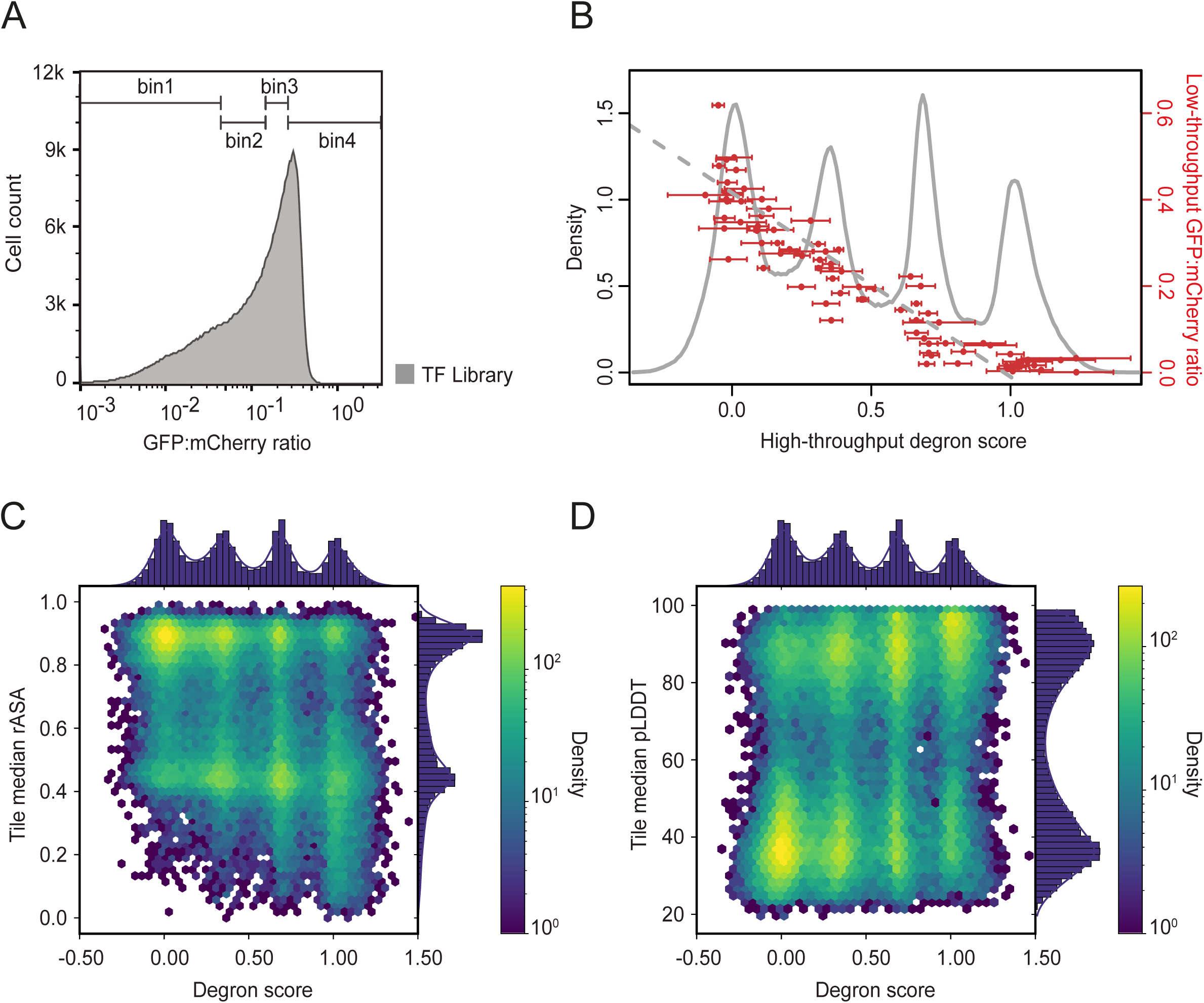
Degrons are mainly found in buried and structured regions. (A) Representative FACS (n=4.84×10^5^) profile displaying the quantification of the GFP:mCherry ratio of cells expressing the TF odd sublibrary. The employed gates (bin 1-4), each containing 25% of the cell population, are marked. (B) Overlayed plot displaying the degron scores of the TF library and the population density (run. avg. size 5) (grey), with the degron scores of 82 randomly selected tiles and the GFP:mCherry ratios of the tiles analyzed in low throughput with flow cytometry (red). The error bars on the high throughput degron scores indicate the standard deviation between replicates. The error bars on the low throughput degrons scores indicate the standard error of the mean and are too small to be visible. (C) Correlation plot of the library degron scores and per-tile median relative accessible surface area (rASA) (absolute Pearson |r|: 0.41, absolute Spearman |*ρ|*: 0.38), and (D) predicted local distance difference test (pLDDT) score (absolute Pearson |r|: 0.36, absolute Spearman |*ρ|*: 0.36) both derived from AF2 models. Yellow indicates high density, and dark blue indicates low density.

### Degrons are mainly located in structured and buried regions

To investigate the overall characteristics of the degrons, identified in this screen, we first examined their positioning within the full-length protein structures predicted by AlphaFold2 (AF2). When comparing the degron scores with the per-tile median relative accessible surface area (rASA) (**Fig. 2C**), we observed that, in general, tiles with a high rASA score, indicating high exposure, displayed low degron scores. Conversely, many tiles with low rASA scores, indicating low exposure, had high degron scores. Hence, the majority of degrons in the TF proteome are not surface exposed, consistent with the assumption that most of the identified degrons are PQC degrons that only operate upon misfolding or unfolding of the protein (44). However, many exposed tiles clearly scored as degrons, which may represent degrons involved in regulated degradation of the native proteins. Of the 21,970 degron tiles (degron score >0.5) with rASA and pLDDT values assigned from AlphaFold models, 6,992 (32%) have rASA >0.7. Previously, it has been suggested that some types of degrons are often found in protein regions that are intrinsically disordered (45). To test this, the degron scores were compared with the median of predicted local distance difference test (pLDDT) score of the tiles, derived from the AF2 predicted structures (**Fig. 2D**). High pLDDT scores indicate high confidence AF2 structure prediction, while tiles with low pLDDT scores indicate low confidence in the structure prediction and disorder. In general, degrons are most commonly found in structured regions (high pLDDT) (**Fig. 2D**), likely indicating that these are PQC degrons. However, we note that many regions with high degron activity are found in regions predicted to be intrinsically disordered (low pLDDT). Thus, of 21,970 degrons (degron score >0.5), 8,817 (40%) have pLDDT <70.

### Exposed degrons regulate the abundance of transcription factors

To investigate the effect of degrons that are surface exposed in the context of full-length proteins and therefore should be constitutively active, we selected two TFs, the pioneer factor achaete-scute homolog 1 (ASCL1, UniProt ID: P50553), known to be constitutively degraded in premature neuronal stem cells (46), and heat shock factor protein 1 (HSF1, UniProt ID: Q00613). These proteins contain tiles with high degron scores that also display high rASA and low pLDDT scores (**Fig. 3AD)**. The proteins were produced as GFP fusions using the same expression system as for the screen (**Fig. 1A**) enabling the abundance of the full-length TFs to be monitored as the relative GFP:mCherry signal by flow cytometry. As expected, we observed a low steady-state level of ASCL1 (**Fig. 3B**) and HSF1 (**Fig. 3E**). The low abundance was due to proteasomal degradation, since in both cases the level increased in response to the proteasome inhibitor bortezomib (**Fig. 3BE**). Deleting part of the ASCL1 degron located within the second tile (position 23-40) in a region predicted to be exposed and disordered (**Fig. 3AC**), resulted in a 7-fold increase in GFP:mCherry ratio (**Fig. 3B**), indicating that indeed this region operates as a degron in ASCL1. However, since the ASCL1Δdegron level was still lower than that observed with proteasome inhibition, other degrons likely also contribute to ASCL1 degradation. Similarly, deletion of part of tile 14 in HSF1 (position 198-217) (**Fig. 3DF**) also led to an increased level of the protein (**Fig. 3E**), suggesting that this region, like the one in ASCL1, contributes to determining the abundance of the TF.

**Figure 3.**
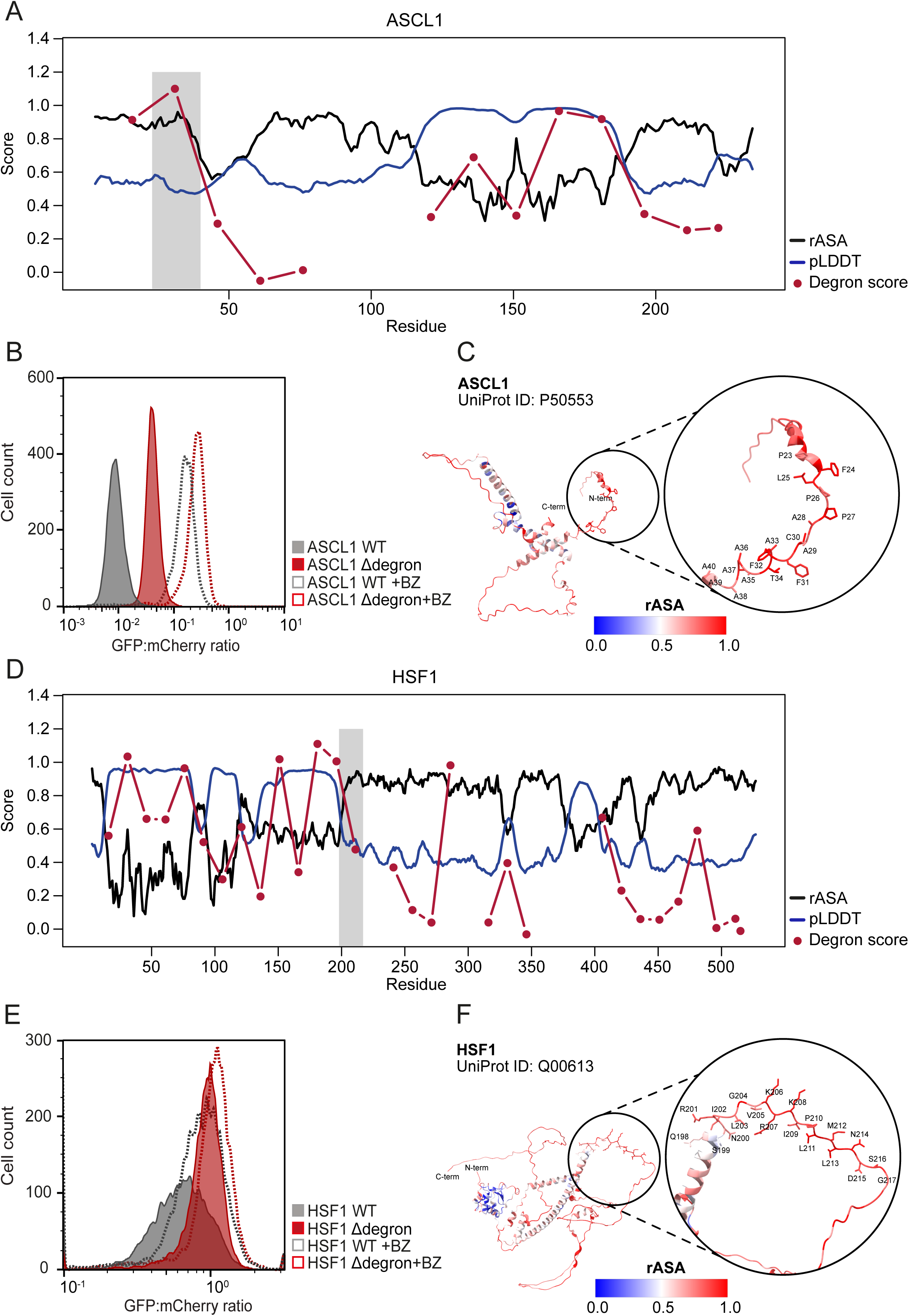
Exposed degrons control transcription factor abundance. (A) Profile of ASCL1 (UniProt ID: P50553) displaying the residue number on the x-axis and per-residue relative accessible surface area (rASA) score (black), predicted local difference test (pLDDT) score (blue) and the degron score (red) on the y-axis (run. avg. size 5). The tile analyzed in panels B and C is shaded. (B) Representative flow cytometry profiles displaying the quantification of the GFP:mCherry ratios of cells expressing wild-type (WT) ASCL1 (n=9,095) and ASCL1 with deletion of amino acids 23-40 (ASCL1Δdegron) (n=9,423), either untreated or treated with 15 μM bortezomib (BZ) for 16 hours (ASCL1 WT + BZ: n=10,280, ASCL1Δdegron + BZ: n=10,669). (C) Full-length protein structure of ASCL1, with the exposed degron highlighted, predicted by AF2 colored in accordance with the per-residue rASA. Blue indicates low rASA and red indicates high rASA. (D) Profile of HSF1 (UniProt ID: Q00613) displaying the residue number on the x-axis and rASA (black), pLDDT (blue) and degron score (red) on the y-axis (run. avg. size 5). The tile analyzed in panels E and F is shaded. (E) Representative histograms displaying the quantification of the GFP:mCherry ratios of cells expressing wild-type HSF1 (n=9,865) and HSF1 with deletion of amino acids 198-217 (HSF1Δdegron) (n=10,355), either untreated or treated with 15 μM bortezomib (BZ) for 16 hours (HSF1 WT + BZ: n=14,669, HSF1Δdegron: n=12,457). (F) Full-length protein structure of HSF1, with the exposed degron highlighted, predicted by AF2 colored in accordance with the per-residue rASA. Blue indicates low rASA and red indicates high rASA.

### Amino acid composition affects degron activity

Next, we used the degron scores, calculated for tiles with a specific amino acid at a defined position, to determine the average effect of that amino acid on the overall stability of the tile. This was visualized as a heatmap of the effect of all possible amino acids at every position in the 30-residue tile (**Fig. 4A**). As previously observed (20, 21, 23, 30, 37), hydrophobic residues in general correlate with high degron scores (reduced protein abundance), while negatively charged residues, and to a lesser extent alanine, glycine, lysine, proline and serine were correlated with low degron scores. The remaining amino acids appeared more or less neutral. These effects are consistent with previous studies on cytosolic proteins in mammalian and yeast cells (20, 21, 23, 30), and therefore suggest that the overall pattern of degron activity between these compartments and cell types is similar. We also note that the enrichment for hydrophobic residues in degrons is in accordance with the concept that many degrons are found in buried and structured regions (**Fig. 2CD**), suggesting that these operate as PQC degrons.

**Figure 4.**
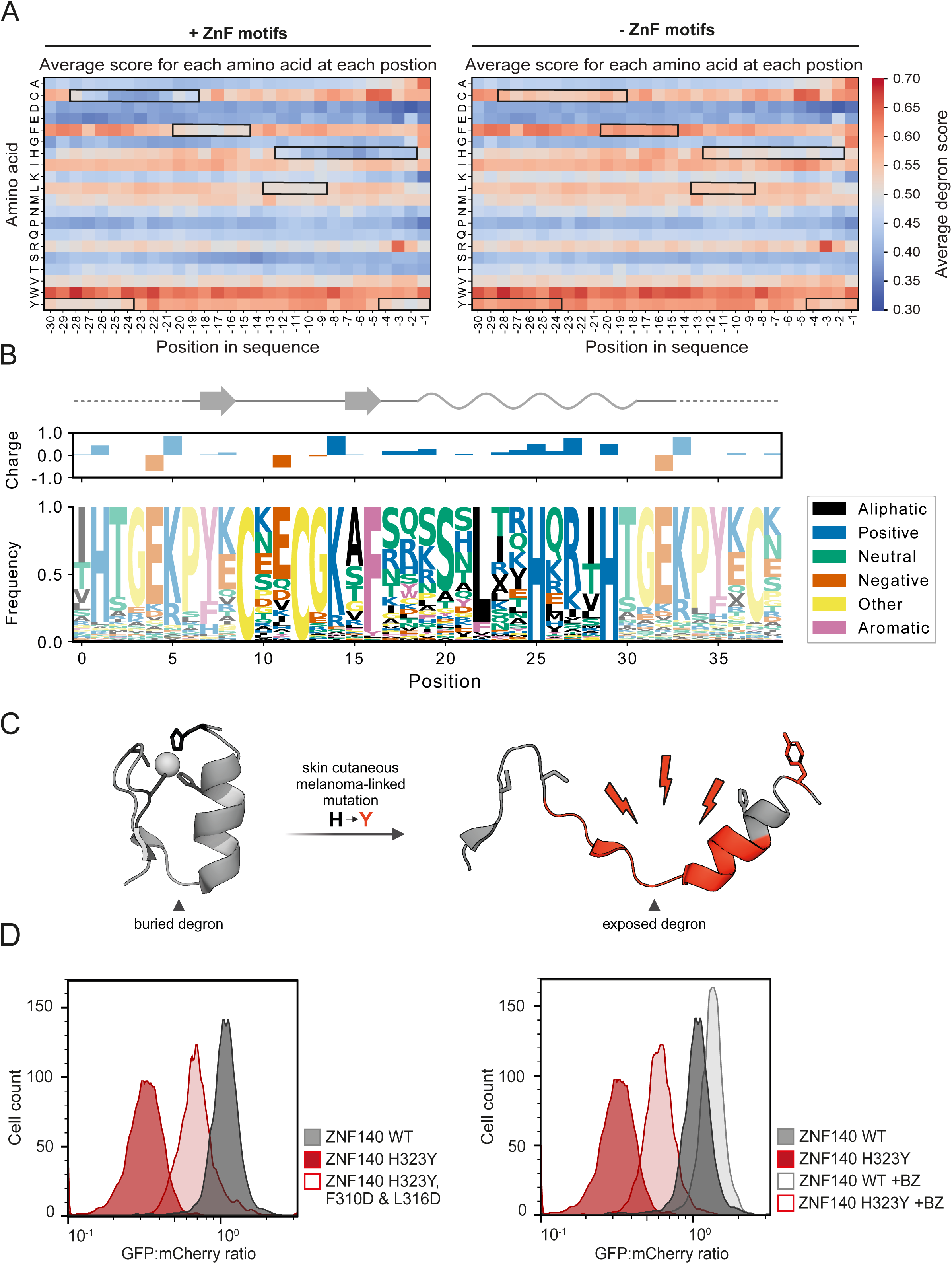
Disruption of the C2H2 ZnF fold leads to exposure of a buried degron. (A) Heatmaps displaying the average degron score of tiles with a specific amino acid at a defined position either with (left panel) or without (right panel) tiles including intact C2H2 ZnF motifs. Red indicates a high degron score and blue indicates low degron score. Stable patches associated with the C2H2 ZnF motifs are framed. (B) Logo plot of all intact C2H2 ZnF motifs (4,282 tiles) extracted from the human transcription factor library colored in accordance with the amino acid properties (see legend). The weighted average charge, based on the frequency of the individual residues at each position is shown, along with the secondary structure elements extracted from the C2H2 ZnF motif at position 378-416 in ZNF140 (UniProt ID: P52738). (C) Schematic cartoon illustrating that mutations linked to skin cutaneous melanoma may lead to disruption of the Zn coordination, unfolding and exposure of a degron. The figure was created with PyMol and BioRender.com. (D) Representative flow cytometry profiles displaying the quantification of the GFP:mCherry ratios of cells expressing tile 21 of ZNF140 (WT) (n=5,000) or the variants H323Y (n=5,000), ZNF140 H323Y, F310D & L316D (n=5,000) either untreated or treated for 16 hours with 15 μM bortezomib (BZ) (ZNF140 WT + BZ: n=5,000, ZNF140 H323Y + BZ: n=5,000).

Apart from the five most C-terminal residues, the degron activity of the tiles appeared largely independent of the amino acid positioning within the tiles (**Fig. S6**), suggesting that, generally, degron potency is determined by the amino acid composition rather than specific sequence. For the five most C-terminal positions, we observe a destabilizing effect of a C-terminal glycine, as well as arginine at the -3 position (**Fig. 4A**), which have previously been described as C-degrons, recognized by specific CRLs (20, 21, 36). A stabilizing effect of a C-terminal lysine residue was also evident (37). Additionally, we observe a destabilizing effect of alanine at position -1 and -2, and cysteines at positions -1 to -5, with the strongest effects seen at positions -4 and -5, which have been attributed to ubiquitin-independent C-degrons (8).

### Disruption of C2H2 Zn finger motif exposes a buried PQC degron

In addition to the C-terminal effects described above, we noted two highly stable clusters of the otherwise destabilizing cysteine and histidine residues, along with regions with leucine, phenylalanine, and tyrosine, which led to increased abundance (**Fig. 4A**). To our knowledge, this pattern has not been reported in previous studies and appears to be unique to our nuclear TF library. To probe this region-specific effect of cysteine, histidine, leucine, phenylalanine and tyrosine residues further (**Fig. 4A**), we first compared the degron activity of all tiles with a fixed cysteine residue at position -25 (**Fig. S7A**). This revealed a stabilizing pattern corresponding to CX_2_CX_12_HX_3_H, where X symbolizes any amino acid residue (**Fig. S7**). This pattern was also evident when analyzing tiles with a fixed cysteine residue at positions -24 and -23 (**Fig. S7BC**). The observed pattern bears resemblance to the Zn finger motif of the C2H2 Zn finger family (47), indicating that potentially folded C2H2 Zn fingers lead to structural stabilization and burial of degrons in the tiles. To test this idea, we identified 690 TFs in our library that contain at least one C2H2 Zn finger motif (from here on termed ZnF); as many TFs contain multiple ZnFs, this led to a total of 6,136 ZnF motifs. From our screen, after tiling, which may disrupt some ZnF motifs, the dataset contains 4,282 intact ZnFs, equivalent to 6.8% of the tiled library. To account for this, we generated a new heatmap (**Fig. 4A**) excluding all intact ZnFs. Indeed, this completely eliminated the stabilizing stretches for cysteine, histidine, leucine, phenylalanine and tyrosine, indicating that the apparent stabilizing effect of these residues is connected with the ZnF regions.

To gain further insight into the nature of the stability connected with the ZnFs, all tiles containing an intact canonical CX_2_CX_12_HX_3_H C2H2 ZnF motif were extracted. The sequences were then aligned by anchoring the conserved cysteine and histidine residues in the motif to create a logo-plot (**Fig. 4B**). From the logo-plot, a strong conservation of the glycine and the lysine at positions +4 and +5 from the first Zn-coordinating cysteine was evident. Additionally, we note that the region between the pairs of Zn-coordinating residues appears to be depleted for negatively charged residues (**Fig. 4B**), but almost invariably contains phenylalanine and leucine residues at positions +7 and +13 after the first cysteine (**Fig. 4B**). The hydrophobic nature of these residues, coupled with the lack of negatively charged residues, could indicate that the region between the C2 and H2 residues contains a PQC degron that is buried by the ZnF fold. Accordingly, mapping the ZnF motif-containing tiles onto the AF2-predicted ZnF structures, revealed rASA and pLDDT scores suggesting very little accessible surface area and high confidence structure (**Fig. S8**). This indicates that a tile of 30 residues is sufficient to drive Zn binding and folding of the small ZnF motif, thus potentially shielding a degron and inhibiting degradation of the tiles.

Previously, it has been shown that substitution of the most C-terminal histidine in several C2H2-type ZnFs to a tyrosine, thus generating a C2HY constellation, is a common somatic mutation linked to skin cutaneous melanoma (40). We hypothesized that such a mutation would reduce Zn interaction and destabilize the ZnF structure, which in turn could lead to exposure of a degron within the hydrophobic loop (**Fig. 4C**). To test this, we selected a tile containing a ZnF from ZNF140 (UniProt ID: P52738) where the H323Y missense variant has been associated with melanoma (40). As above, the tiles were produced as recombinant GFP fusions, and the amounts were determined by flow cytometry. Indeed, the melanoma-linked H323Y missense variant displayed clearly reduced levels compared to the wild-type (WT) (**Fig. 4D**). The reduced abundance was due to proteasomal degradation, since the level was increased by treatment with bortezomib (**Fig. 4D**). Since the low level was independent of whether H323 was replaced with a tyrosine or an alanine residue, and which of the Zn-coordinating residues that were mutated (**Fig. S9**), the reduced abundance is likely due to disruption of the Zn-coordination, leading to unfolding and degron exposure. Accordingly, replacing the conserved phenylalanine and leucine residues with acidic residues increased the abundance of the H323Y variant (**Fig. 4D**).

Collectively, these results indicate that disruption of any of the Zn-coordinating residues in ZnF motifs leads to unfolding, degron exposure, and proteasomal degradation, providing a mechanistic basis for the melanoma-linked H323Y missense mutation.

### Different C-degron characteristics define degron activity in the cytosol and nucleus

Recently, we mapped degrons within the cytosolic human proteome using a similar experimental approach in the same cell line (37). Hence, for the subset of proteins overlapping between the two libraries, it is possible to test if differences in degron composition make them more or less susceptible to degradation in the cytosol or nucleus. To this end, tiles that were common amongst the two libraries were identified, resulting in a list of 234 proteins corresponding to 8,645 tiles. Mapping the degron scores of the tiles from the nuclear screen to the cytosolic screen (**Fig. S10**), as well as calculated abundance scores (**Fig. 5A**), revealed an overall high correlation between the two datasets (Pearson correlation 0.95 and 0.92, respectively) (**Fig. 5A**). The abundance score is a non-linear re-normalized version of the degron score applied to handle the four-peak distribution and ease the comparison of experiments (see Methods). In addition, many of the TF degrons could be identified using the computational peptide abundance predictor (PAP) (37) trained on a dataset of tiles produced in the cytosol (**Fig. 5B**).

**Figure 5.**
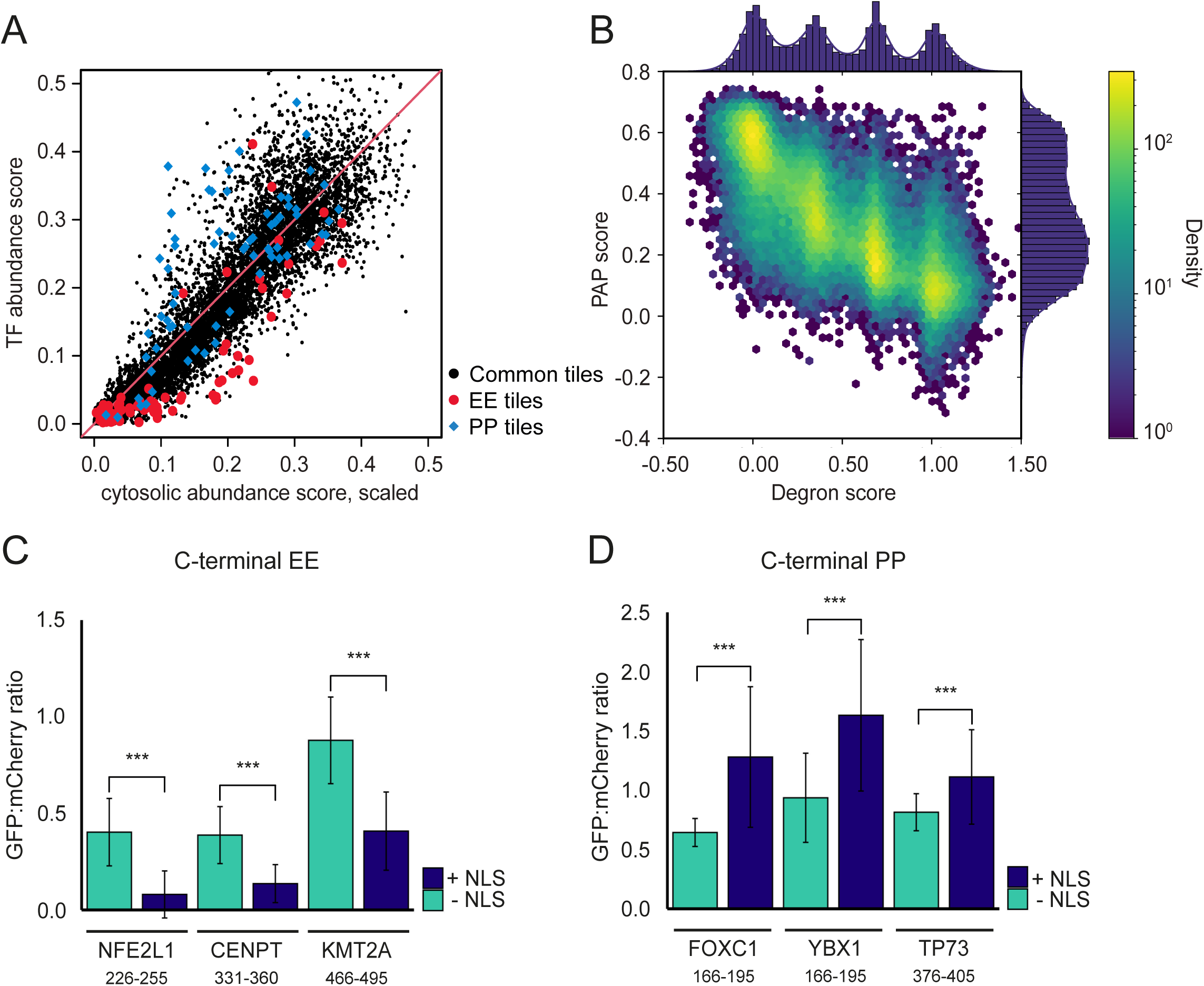
Different characteristics determine degron activity in the nucleus compared to the cytosol. (A) Correlation plot with calculated abundance scores of common tiles between the human transcription factor library and the human cytosolic library (8,645 tiles) (37) (absolute Pearson |r|: 0.92, absolute Spearman |*ρ|*: 0.95). All common tiles are marked with black dots. Tiles ending in EE found to have significantly lower abundance in the nucleus (p < 10^-6^) are marked with red dots. Tiles ending in PP found to have significantly lower abundance in the cytosol (p < 10^-3^) are marked with blue diamond. (B) Correlation plot of the human transcription factor degron scores and peptide abundance predictor (PAP) scores (absolute Pearson |r|: 0.79, absolute Spearman |*ρ|*: 0.80). Yellow indicates high density, and dark blue indicates low density. (C) Bar plot illustrating the difference in mean GFP:mCherry ratio of HEK293T cells expressing tiles ending in EE from the proteins NFE2L1 (residues 226-255), CENPT (residues 331-360) and KMT2A (residues 466-495) either with or without an NLS (*** = p < 0.001, Welch’s t-test). (D) Bar plot illustrating the difference in mean GFP:mCherry ratio of HEK293T cells expressing tiles ending in PP from the proteins FOXC1 (residues 166-195), YBX1 (residues 166-195) and TP73 (residues 376-405) either with or without an NLS (*** = p < 0.001, Welch’s t-test).

The above data suggests that most degrons are equally functional in the cytosol and nucleus. However, this does not rule out the possibility that some rare degrons may be differentially targeted. To identify such degrons, we next turned our attention to the outliers in the correlation of the calculated abundance scores between the cytosolic- and nuclear-expressed tiles (**Fig. 5A**). Among these, several tiles ended with EE or PP (**Fig. 5A**). A paired two-sided *t*-test showed that tiles ending in PP are significantly more abundant in the nucleus compared to the cytosol (*p* < 10^-3^) (**Fig. 5A**). In contrast, tiles ending in EE are significantly more abundant in the cytosol compared to the nucleus (*p* < 10^-6^) (**Fig. 5A**), suggesting that this C-degron is stronger in the nucleus. To test this, we performed low throughput flow cytometry analyses on selected tiles expressed with or without an NLS. For normalization, we included a tile present in both libraries (with an abundance on the diagonal in **Fig. 5A**). The results confirmed that tiles with the EE C-degron are indeed less abundant in the nucleus (**Fig. 5C**), while tiles with a C-terminal PP are less abundant in the cytosol compared to the nucleus (**Fig. 5D**). These results indicate that while most degrons exhibit similar degron potency between the nucleus and the cytosol there are exceptions.

### Transcription factor activity partially explained by protein abundance levels

Recently, a high throughput deep mutational scan was performed to map variant effects on transactivating activity of the cone-rod homeobox (CRX) TF (48), which AF2 predicts is largely disordered (high pLDDT scores) except for a structured homeodomain (residues 40–95) (**Fig. S11A**). Since the read-out of this assay was not normalized to protein abundance levels, the non-functional variants possibly include loss of abundance variants, while variants with an increased activity could include high abundance variants. Thus, to investigate a possible overlap in changes in transactivation and protein abundance, we predicted the change in degron potency of all investigated CRX variants using PAP and the protein language model, ESM-1b, known to efficiently predict variant effects, though with reduced accuracy for disordered regions (49, 50). The results revealed that variant effects in the IDRs of CRX are better explained by ΔPAP scores compared to ESM-1b scores (Pearson 0.54 vs 0.31 for ΔPAP and ESM-1b, respectively) (**Fig. S11BCD**). Conversely, in the folded homeodomain, the mutational effects correlated better with ESM-1b scores compared to ΔPAP scores (Pearson 0.52 vs. -0.19 for EMS-1b and ΔPAP, respectively). This suggests that an alteration of the degron potency can provide a mechanism for variant effects in IDRs.

### Degron properties of transactivating motifs

Previously, it has been suggested that transcriptional activation domains (TADs) and degrons overlap (4). To test this systematically, we utilized ADpred (51) to identify tiles containing activation domains and compared their degron scores with the entire TF library (see methods). Indeed, this revealed that many tiles containing TADs display high degron scores (**Fig. 6A**), suggesting that they play an additional role in affecting TF degradation. Of 1,570 TAD tiles (ADpred >0.8), 1,515 (97%) have measured degron scores, and 1,017 (67%) score >0.5 and 407 (27%) score >1.0. In agreement with this, many TADs were predicted by PAP to display a reduced abundance (**Fig. 6B**), *i.e*. of 1,570 TAD tiles with PAP scores, 892 (57%) have scores <0.2.

**Figure 6.**
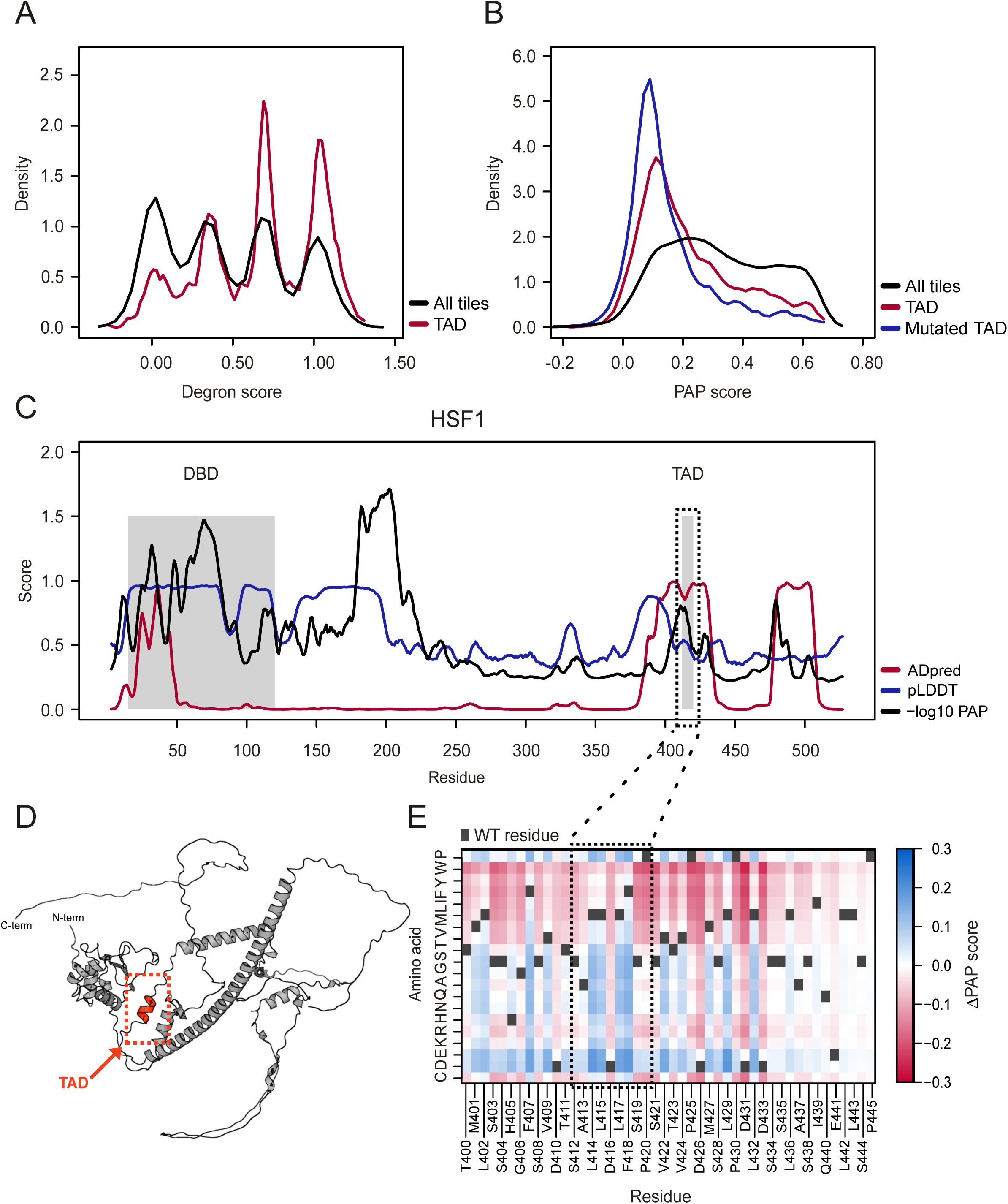
Acidic residues in transactivating domains reduce degron potency. (A) Degron score distributions of the TF library (black) and the tiles predicted to contain a transactivation domain (TAD) (1570 tiles) using ADpred (51) (red) (run. avg. size 5). (B) Distributions of peptide abundance predictor (PAP) scores of all tiles in the TF library (black), tiles predicted to contain a TAD using ADpred (red) and tiles predicted to contain a TAD where the most central acidic residue (D or E) is mutated to a valine (blue) (run. avg. size 5). (C) Profile of HSF1 (UniProt ID: Q00613) displaying the residue number on the x-axis and the -log_10_ PAP score (black), predicted local difference test (pLDDT) score (blue) and the ADpred score (red) on the y-axis (run. avg. size 5). The DNA binding domain (DBD; residues 15-120) and the TAD domain are highlighted by grey boxes. (D) Full-length protein structure of HSF1 (UniProt ID: Q00613) predicted by AF2. The TAD at position 412-420 is marked in red. (E) A PAP-map corresponding to the area marked on the HSF1 profile (panel C) illustrating the calculated ΔPAP scores of all single amino acid substitutions. Blue indicates high protein abundance and red indicates low protein abundance. The WT residues are marked in black. A running average of five residues was applied to all distributions and profiles in this figure.

In general, transactivating regions are enriched for hydrophobic and acidic residues (51-53). Since the presence of hydrophobic residues agrees with the TADs operating as degrons, while the presence of acidic residues does not, we speculated that the acidic residues may play a role in protecting some TFs from degradation (38). To examine this, we used PAP to predict how substituting a single D or E to a V in a TAD, and found that these were predicted to reduce the protein abundances of essentially all TADs (**Fig. 6B**). To further illustrate this, we selected HSF1 and used ADpred to map TADs in this transcription factor. HSF1 contains a DNA binding domain (DBD) in the N-terminal region, while the ∼300 C-terminal residues are predicted to be disordered (low pLDDT) (**Fig. 6C**). ADpred identifies two TAD regions towards the C-terminus, including the previously noted (54) 412–420 TAD (**Fig. 6C**), which is predicted to form an exposed helix (**Fig. 6D**). As a proxy for degron potency, we overlayed -log(PAP) predictions with the HSF1 ADpred scores (**Fig. 6C**), revealing a distinct peak of increased predicted degron activity within the 412–420 TAD. Next, ΔPAP scores for all possible single amino acid substitutions in HSF1 (**Supplemental Data File**) were plotted as a heatmap for the TAD region (**Fig. 6E**). This showed that mutation of the acidic residue D416 residue to any non-acidic residue is predicted to reduce abundance, while mutation of the hydrophobic L414, L415, L417 and F418 residues to hydrophilic residues is predicted to increase abundance (**Fig. 6E**). This example and the analysis on all TADs (**Fig. 6AB**), suggests that the hydrophobic residues contribute to the degron potential of TADs, while the acidic residues counteract this effect.

## Discussion

As key regulatory proteins, many TFs are subject to rapid degradation (4). Here, we comprehensively mapped degrons in the entire human TF proteome, revealing both exposed and buried degrons specifically determining the abundance of TFs in a proteasome-dependent manner. We find that the degron potential between tiles in the nucleus and cytosol generally correlate, but certain sequence motifs affect cytosolic and nuclear abundance differently. Thus, prediction of protein abundance, using the PAP machine-learning model trained on cytosolic proteins (37), match our experimental data on nuclear TFs. This indicates that most degrons are functional in both the nucleus and the cytosol. However, since different components of the degradation machinery, such as E3s and chaperones, may be differentially localized, some differences in degron recognition are expected. DCAF12, the adaptor protein of CRL4, which is responsible for substrate recognition of glutamate in position -2 (39) has two N-terminal NLSs and forms unstable complexes in the cytosol, where it is rapidly degraded (28). This could explain the fact that the EE-ending tiles are less abundant in the nucleus compared to the cytosol. Similarly, proline-ending peptides are targeted by FEM1b (39), a CRL2 adaptor protein. FEM1b is predominantly localized in the cytoplasm (55), which could explain the relatively lower abundance of PP-ending tiles in the cytosolic library. Possibly there are many more examples such as these. However, they are not readily identifiable in our dataset, since only a limited number of tiles were tested in both compartments.

Recent studies have demonstrated that various nuclear proteins, including transcription factors, encoded by the immediate-early genes, are degraded by the proteasome independently of ubiquitin, but instead reliant on the proteasomal co-factor midnolin (7). Using its Catch domain, midnolin engages disordered regions in its substrates that upon binding form a β-strand within the hydrophobic core of the Catch domain. Presumably, several of the degrons identified here are degraded via this pathway. However, as we note that most of the identified degrons are stabilized upon inhibition of the ubiquitin E1, ubiquitin-dependent degradation appears to be the preferred route for our tiled library.

Similar to other proteins (20, 22, 30), the main feature of degrons in TFs appear to be hydrophobicity. Since such regions tend to be buried, most of the identified degrons are located in the structured DNA binding domains and thus likely function as PQC degrons. As shown for the H323Y cancer-linked somatic missense mutation in ZNF140 (40), this likely results in a dramatic destabilization of the ZnF fold, resulting in exposure of the non-charged and hydrophobic loop between the pairs of Zn coordinating residues, thus allowing this region to operate as a degron. We suspect that many of the identified PQC degrons may similarly trigger the degradation of TFs that become structurally destabilized due to mutations or environmental stress conditions.

Although most of the identified degrons are located in the structured parts of the TFs, we do find several within the intrinsically disordered regions. As shown for ASCL1 and HSF1, these appear to be active also in the context of the native full-length proteins and thus contribute to keeping the steady-steady state level of these TFs low. As a pioneer TF involved in cell differentiation (56) it is likely critical that the level of ASCL1 is kept low. Similarly, in the case of HSF1, which drives the expression of stress response genes (57), unwarranted expression is likely to have adverse effects. Indeed, constitutive HSF1 transcriptional induction is known to inhibit cell growth (58). That predictable degradation signals are constitutively active in IDRs and important for TF function is supported by a good correlation between the CRX transactivation assay and PAP scores, highlighting also that it may be difficult to disentangle effects on abundance and intrinsic function.

With the present screening approach where the TFs are tiled into shorter peptides, it is unlikely that the tiles are subject to substantial post-translational modifications (PTMs). Accordingly, we are probably limited to identifying constitutive degrons, which do not depend on PTMs for activation. Thus, phospho-degrons, which rely on prior phosphorylation, are likely not detected in our screen. In addition, since negatively charged residues appear to lower the degron activity, it is possible that some of the exposed degrons are blocked by phosphorylation in the full-length context and would thus require dephosphorylation to become active.

In addition to the folded DNA binding domains, most TFs also contain long IDRs in which the TADs typically reside (59). Using a similar tiling approach as the one we applied here, recent high-throughput studies have aimed at mapping the motifs involved in transactivation (53, 60). This revealed a surprisingly large number of sequences with transactivating function. In general, TADs are rich in hydrophobic residues but also contain acidic residues (52, 53). An overlap between TADs and degrons has been noted for years and has been suggested as a mechanism to ensure degradation of active TFs (4). According to the results presented here, the hydrophobic residues in TADs are consistent with them operating as degrons, while the presence of acidic residues is not. We speculate that the acidic residues may play a dual role including preventing TA regions from being recognized as degrons, which in turn suggests that during evolution, the UPS may have contributed to shape the TADs. Degrons doubling as TADs can also have a direct consequence for interpreting the results from large-scale studies on mapping transactivating motifs, since it is unknown how the transactivating activity in e.g. a yeast 1-hybrid setting depends on the abundance of the hybrid TF protein, and it may be difficult to correct for such abundance effects. The results presented here may be useful to account for TF abundance in such assays, but also for a general functional and mechanistic characterization of TFs and for predicting effects of disease-linked coding variants in TFs.

## Methods

### Library design

The TF library was designed based on the catalogue of 1639 human transcription factors described by Lambert *et al*. (41). For each gene, a nucleotide sequence was obtained using the MANE Select list of transcripts version 1.0 (61) and Ensemble GRCh38 protein coding sequences release-107 (both downloaded May 2022). For 13 genes not in the MANE Select list, we used the Ensembl canonical transcript if this was annotated with experimental evidence at protein level. For two genes, DUX1 and DUX3, sequences were obtained from European Nucleotide Archive. A total of 1626 protein coding sequences were obtained after filtering 11 depreciated entries and two synonymous sequences. The library was designed as 90 nt (30 residues) tiles overlapping with 45 nt (15 residues) and split into three non-overlapping sublibraries of 30,292 even, 31,080 odd and 1,620 C-terminal tiles, which are each unique at the nucleotide level. Nucleotide sequences contained a stop codon causing the C-terminal tiles to be only 29 amino acids long. In total, the library consists of 63,230 tiles of which 62,992/62,808 are unique at nucleotide/amino acid level (excl. control peptides), respectively.

### Plasmids

The TF library was cloned into an integrative mammalian expression vector directly fused to GFP containing an N-terminal NLS (42). Variants of ASCL1, HSF1, and ZNF140 were introduced into the same mammalian expression vector as described above (Genscript). Deletions and point mutations were generated by Genscript.

### Cloning

The TF libraries were produced by Twist Biosciences. To reduce the probability of template switching during PCR, the TF library was divided into 3 libraries, the evens (complexity: 30,292), odds (complexity: 31,080) and C-terminal (complexity: 1,620) libraries. The oligo libraries were designed with two adaptors that included complementary sequences (5’ complementary sequence: GTTCTAGAGGCAGCGGAGCCACC, 3’ complementary sequence: TAGTAACTTAAGAATTCACCGGTCTGACCT) to the flanking regions of the p0489_AttB_NLS-EGFP-Link-IRES-mCherry_Bgl2 expression vector. In order to compare scores between the different sublibraries, we supplemented each sublibrary with five synonymous iterations of three control tiles as well as one iteration of ten control tiles, for a total of 25 control tiles ranging from 66 – 90 nt in length. The control tiles were selected based on their degron scores determined in previous studies (62, 63).

The libraries were dissolved in nuclease-free water (Thermo Fisher Scientific, AM9937) to a final concentration of 20 ng/µL (odds and evens) and 10 ng/µL (C-terminal). The libraries were amplified into double-stranded DNA oligos in a 50 µL reaction using 10 ng of the even and odd, and 5 ng of the C-terminal library, respectively, with 0.5 µM forward primer VV3 and reverse primer VV4 and using 25 µL Q5 High-Fidelity 2x Master Mix (New England Biolabs, M0492L). The oligoes were denatured at 98 °C for 30 sec, followed by 8 PCR cycles: denaturing for 5 sec at 98 °C, annealing for 30 sec at 69 °C, elongation for 10 sec at 72 °C, with a final extension at for 2 min at 72 °C. The reaction products were purified by separation on a 2% agarose gel using 1x SYBR Safe DNA Gel Stain (Thermo Fisher Scientific, S33102) followed by gel extraction using GeneJet Gel Extraction Kit (Thermo scientific, K0691) with elution in 30 µL nuclease-free water.

The expression plasmid, p0489_AttB_NLS-EGFP-Link-IRES-mCherry_Bgl2, was amplified and linearized in a 50 µL reaction using 5 ng plasmid DNA, 0.5 µM forward primer VV1 and reverse primer VV2, and 25 µL Q5 High-Fidelity 2x Master Mix. The PCR reaction was run as the following: 98 °C for 30 sec, followed by 21 cycles of: 98 °C for 5 sec, 69 °C for 30 sec, 72 °C for 3 min and 40 sec, with a final extension for 2 min at 72 °C. Reaction products were cleaned and concentrated using DNA Clean & Concentrator-5 (Zymo Research, D4014) with elution in 10 µL nuclease-free water. Methylated DNA was removed in a 50 µL reaction using the concentrated 10 µL vector DNA, 2 µL DpnI (New England Biolabs, R0176L) and 5 µL 10x CutSmart buffer (New England Biolabs, B7204). The reaction was incubated overnight at 37 °C, followed by a 2-hour incubation with 2 µL additional DpnI. Subsequently, the reaction product was purified on a 1% agarose gel using 1x SYBR Safe DNA Gel Stain, followed by gel extraction using GeneJet Gel Extraction Kit (Thermo scientific, K0691) and elution in 30 µL nuclease-free water. The Qubit 1X dsDNA High Sensitivity (HA) kit (Thermo Fisher Scientific) was used to determine the DNA concentration.

The tiled TF library oligoes were cloned into the expression vector using Gibson Assembly (64). Gibson assembly was performed in a 20 µL reaction with a vector to oligo molar ratio of 1:4 using 10 µL 2x Gibson Assembly Master Mix (New England Biolabs, E2611L) incubated at 50 °C for 1 hour. Reaction products were cleaned and concentrated using DNA Clean & Concentrator-5 eluting in 12 µL nuclease-free water.

For bacterial transformation, 50 µL NEB10β electro competent *E. coli* cells (New England Biolabs, C3019I) were thawed on wet ice. Once thawed, 2 µL Gibson assembly reaction product was transferred to the vial followed by electroporation (2 kV). Cells were resuspended in 950 µL pre-warmed NEB10β Outgrowth Medium (New England Biolabs, B9035S), followed by incubation at 37 °C for 1 hour in a shaking incubator. To maintain full coverage of the TF library, five individual transformations of each library were performed. From each transformation, 20 µL were diluted 1:1000 in SOC media and 100 μL was plated on LB agar plates containing 100 µg/mL ampicillin. Plates were incubated overnight at 37 °C and colonies were counted to ensure at least 100-fold coverage of the complexity of each library. The remaining 980 μL of each transformation was transferred to the same 200 mL LB media containing ampicillin. The flask was incubated with shaking overnight at 37 °C. The cells were harvested by centrifugation at 3000 g for 10 min and plasmids were purified using NucleoBond Xtra Midi kit (Macherey-Nagel, 740410.50). The DNA yield and purity of the purified plasmids were measured by a NanoDrop spectrometer ND-1000.

Tiles investigated individually in low throughput were synthesized by Integrated DNA Technologies (IDT). The individual tiles were cloned into the p0489_AttB_NLS-EGFP-Link-IRES-mCherry_Bgl2 or the p2127_attB-EGFP-PTEN-IRES-mCherry-562bgl by Gibson assembly following the protocol described above. The tiles used for low throughput validation of the degron scores were randomly picked from transformed plates from the three sublibraries and also include 31 tiles covering PPARγ (UniProt ID: E9PFV2) that were cloned individually.

### Cell culture

The HEK293T landing-pad cell line TetBxb1BFPiCasp9 Clone 12 (43) was cultured in high glucose Dulbecco’s Modified Eagles Medium (Sigma-Aldrich, D5796) supplemented with 10% fetal bovine serum (FBS) (Sigma-Aldrich, F7524), 0.29 mg/mL penicillin G potassium salt (BioChemica, A1837,0025), 0.24 mg/mL streptomycin sulphate (BioChemica, A1852,0025), 0.32 mg/mL L-glutamine (Sigma-Aldrich, G3126) and 2 μg/mL doxycycline (Sigma Aldrich, D9891), at 37 °C in a humidified incubator with 5% CO_2_. Cells were passaged at ∼80% confluence. Cells were tested negative for mycoplasma (Mycostrip, InvivoGen, ref-mys-10), and identity was regularly confirmed by microscopy as doxycycline-inducible blue fluorescence and by counter selection with 10 nM AP1903 (MedChemExpress, HY-16046). Transfections were performed in media without supplemented doxycycline using Fugene HD (Promega, E2312) with pCAG-NLS-Bxb1 (Addgene, Plasmid #51271) plasmid at a molar ratio of 1:17.5 with the expression plasmid. For transfection in a 15 cm plate (and proportionally less for smaller plates), 19.8 μg plasmid DNA was used in a final volume of 3630 μL optiMEM (Thermo Fisher Scientific, 11058021) and 74.1 μL Fugene HD. The mixture was incubated for 15 min at room temperature and subsequently gently added to the cell media by pipetting. After 48 hours, 10 nM AP1903 (MedChemExpress, HY-16046) was added to fresh doxycycline-containing media to select for successful recombinants. The cells were flow sorted four days after addition of doxycycline and AP1903. A minimum of 4×10^6^ cells was maintained at all times after library recombination to ensure at least 100-fold coverage of the libraries.

### Flow cytometry

For flow cytometry, cells were dislodged using trypsin (Gibco, 27250-018) and pelleted by centrifugation at 300 g for 5 min, followed by resuspension in PBS (10 mM Na_2_HPO_4_, 1.8 mM KH_2_PO_4_, 137 mM NaCl, 3 mM KCl, pH 7.4) with 2% FBS and filtered through a 50 μm polyethylene mesh filter. Subsequently, cells were analyzed with a BD FACSJazz or BD Fortessa (BD Biosciences) instrument using excitation lasers 405 nm, 488 nm and 561 nm for BFP, GFP and mCherry, respectively, and filters 450/50 (FACSJazz) or 431/28 (Fortessa) for BFP, 530/40 (FACSJazz) or 530/30 (Fortessa) for GFP and 610/20 (FACSJazz and Fortessa) for mCherry. Data were collected and analyzed with FlowJo v10.09. Live cells and singlets were gated using forward and side scatter, followed by gating of successfully transfected cells based on BFP negative and mCherry positive events (**Fig. S12**).

Perturbations were performed prior to flow cytometry analysis as follows: 15 μM bortezomib (LC Laboratories, B-1408) for 16 hours, 20 μM chloroquine (Sigma Aldrich, C6628) for 16 hours, 1 μM MLN7243 (MedChemExpress, HY-100487) for 16 hours, 2 μM MLN4924 (MedChemExpress, HY-70062) for 16 hours and 10 μg/ml cycloheximide for 8 hours (Sigma-Aldrich, 01810).

### Cell sorting

Fluorescence-Activated Cell Sorting (FACS) was performed using an ARIA III FACS instrument (BD Biosciences) equipped with a 70 μm nozzle and using the FACSDiva software. The cells were analyzed using excitation lasers 405 nm, 488 nm and 561 nm for BFP, GFP and mCherry, respectively, and filters 442/46, 530/30 and 615/20 for BFP, GFP and mCherry, respectively. Live cells and singlets were gated using forward and side scatter, followed by gating of successfully transfected cells based on BFP negative and mCherry positive events (**Fig. S13**).

Samples were prepared by dislodging of cells using trypsin and centrifugation at 300 g for 5 min, followed by resuspension in PBS with 2% FBS for a concentration of 10×10^6^ cells/mL. Cells were sorted into 4 bins based on their GFP:mCherry ratio ensuring 25% of the population in each bin with a minimum of 4×10^6^ cells for the even and the odd library, and 2×10^5^ cells for the C-terminal library in each bin to ensure full coverage of the libraries. The sorted cells were pelleted by centrifugation at 300 g for 5 min, resuspended in fresh media without doxycycline and transferred to individual cell culture plates. Cells were cultured for an additional four days. After four days, the cells were harvested by centrifugation at 300 g for 5 min, the supernatant was aspirated, and the cell pellet was stored at -80 °C. Three biological replicates (separate transfections) of each library and two sorting replicates were performed.

### Genomic DNA extraction

Genomic DNA was extracted from ∼15×10^6^ sorted cells of each library replicate using DNeasy Blood and Tissue Kit (Qiagen, 69504) following the protocol “Purification of Total DNA from Animal Blood or Cells” except for the following differences: Step 1c: addition of 20 μL 10 mg/mL RNase A instead of 4 μL 100 mg/mL RNase A. Step 2: incubation at 56 °C for 30 min with intermediate shaking instead of incubation at 56 °C for 10 min. Step 4: Centrifugation at ≥ 6000 g for 2 min instead of 1 min. Step 7: Elution in 150 μL nuclease-free water instead of elution in 200 μL Buffer AE/C.

### Genomic DNA amplification, purification and quantification

The tile inserts were isolated from the extracted genomic DNA using adapter PCR. For each bin, ten 50 μL reactions were prepared using 2.5 μg template DNA, 0.5 μM forward primer VV7 and reverse primer VV8 and 25 μL Q5 High-Fidelity 2x Master Mix. Samples were denatured for 30 sec at 98 °C followed by 7 cycles of denaturation for 10 sec at 98 °C, annealing for 30 sec at 65 °C and elongation for 50 sec at 72 °C, and a final extension for 2 min at 72 °C. A technical replicate was performed for each bin. The PCR products were cleaned using Ampure XP bead-based reagent (Beckman Coulter, A63881) (0.8:1 ratio) and eluted in 20 μL nuclease-free water.

The isolated DNA fragments were next shortened and indexed by qPCR in a 50 μL reaction using 4 μL of the Ampure XP bead cleaned PCR product, 0.5 μM of either of the following forward primers; VV29, VV30 or VV31, and 0.5 μM indexing reverse primer JS_R, varying for each sample, 0.5 x SYBR Green Nucleic Acid Gel Stain (Thermo Fisher Scientific, 2311857) and 25 μL High-Fidelity 2x Master Mix. Samples were denatured at 98 °C for 30 sec, followed by 15 cycles of denaturing for 10 sec at 98 °C, annealing for 30 sec at 67 °C, elongation for 15 sec at 72 °C. The PCR products were purified on a 2% agarose gel using 1x SYBR Safe DNA Gel Stain followed by gel extraction using GeneJet Gel Extraction Kit with elution in 30 µL nuclease-free water. The concentration of the samples was quantified using a Qubit 2.0 Fluorometer (Invitrogen) with a Qubit 1X dsDNA High Sensitivity (HS) Kit (Thermo Fisher Scientific, Q33231).

### Sequencing

The libraries were sequenced along with 10% PhiX (Illumina), using a NextSeq 550 sequencer (Illumina) with a NextSeq 500/550 High Output Kit v2.5 (300 Cycles) (Illumina, 20024905) using custom sequencing primers VV17-C and VV25-C for read 1 and read 2 (paired-end), respectively, and primers VV19-C and VV16-C for index 1 and index 2, respectively. The prepared amplicons from each bin were mixed, ensuring at least a 100-fold coverage of their equivalent sublibrary complexity.

### Calculating degron and peptide abundance predictor (PAP) scores

Sequencing reads were processed as in (37). Briefly, sequencing reads with an exact match to the library (or controls) were counted and the protein stability index (PSI) was calculated per tile using:

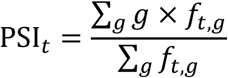

where *f_t,g_* is the frequency of tile *t* in FACS gate *g*. We require 100 or more reads across the four gates. All individual replicates had scores calculated for 96-98% of library members. Each of the three sub-libraries have 6 replicates, 3 biological times 2 FACS replicates, with an average PSI Pearson correlation of 0.97 between all replicate pairs (range 0.91-0.99). One replicate of the CT library failed. As previously, the scores are observed to distribute in four peaks which we consider an artifact of the sort-seq experiment and an indication of a low noise level. A degron score is obtained by normalizing each replicate such that the median of the stable peak has degron score zero and most unstable peak a score of one. The three libraries are combined by averaging normalized scores over replicates. We require two or more replicate measurements per tile and report the standard deviation between replicates as score uncertainties resulting in 58,781 scores which are mapped to 58,992 tiles in the original (redundant) library. An abundance score is obtained by transforming (non-linearly) the degron scores to the fluorescence distribution observed during FACS with error propagation (37). Peptide Abundance Predictor (PAP) scores were calculated using the PAP software package (37) available via GitHub (https://github.com/KULL-Centre/_2025_Voutsinos_degron_cytosol). The C-degron term was incorporated into the prediction for each tile to account for the influence of C-terminal sequence features on peptide stability and degradation. In the evaluation of full-length proteins, the C-degron term is only applied to the C-terminal tile.

### rASA and pLDDT predictions

First, we collected protein structures from version 4 of the AlphaFold2 Protein Structure Database (AFDBv4) (https://ftp.ebi.ac.uk/pub/databases/alphafold/v4/) (65, 66). Proteins shorter than 16 residues, those containing non-standard amino acids, or those absent from the AlphaFold UniProt reference proteome ‘one sequence per gene’ FASTA file were excluded from our analysis. For proteins exceeding 2,700 amino acids in length, as their AFDBv4 structure predictions are segmented into 1,400-amino-acid-fragments with a 1,200-amino-acid-overlap, we calculated exposure for each fragment and extracted the pLDDT values for each fragment. The average of all the values from overlapping fragments at a given residue position was then taken as the representative value for that residue for both metrics.

We extracted the predicted local distance difference test (pLDDT) scores from the AF2 structures, using the per-residue measure of local confidence as a determinant of intrinsically disordered regions (67). The relative Accessible Surface Area (rASA) was calculated using DSSP (68) with maximum solvation areas from (69).

### Sequence logo generation

We identified 4,282 tiles from our TF tile sequence library containing an intact C2H2 ZnF motif, defined by the canonical consensus pattern (CX_2_CX_12_HX_3_H). Each motif was aligned so that the conserved cysteine and histidine residues remained in register, thereby preserving the core of the motif. We then generated a sequence logo from these aligned motifs, with residues colored according to their biochemical properties: positive residues (H, K, R) are shown in blue, negative residues (D, E) in orange, neutral residues (N, Q, S, T) in green, other residues (C, G, P) in yellow, aliphatic residues (A, I, L, M, V) in black, and aromatic residues (F, Y, W) in purple.

### Predicting transactivating regions

Transactivation regions were predicted using the ADpred (51) python module version 1.2.8 with PSIPRED 4.0 features obtained from the PSIPRED workbench API (http://bioinf.cs.ucl.ac.uk/psipred). Predictions were made for full-length proteins. As for this work, ADpred considers tiles of length 30, so we used the score of residue 16 of each tile to represent this. We used default settings including the end-rule of extending the chain with 15 positions of Gly in coil secondary structure in both ends. Tiles are considered transactivating if the score exceeds 0.8, reported to be a conservative threshold by the developers of the method (51).

## Acknowledgements

The authors thank Anne-Marie Bonde Lauridsen, Søren Lindemose, Isa K. Henrichs and Rajesh Somasundaram for excellent technical assistance. We thank Vibe H. Østergaard and Michael Lisby for assistance with the FACSJazz instrument. We also thank Morten Meyer and all members of the PRISM and REPIN centers for helpful discussions. We acknowledge the use of the FACS, sequencing and computing core facilities at the Biotech Research & Innovation Centre and Department of Biology, University of Copenhagen. Fig. 1A and 4C were created with BioRender.com. Fig. 4C and 6D were created with PyMol.

## Supplemental files

This article includes the following supplemental information:

- Supplemental figures (SupplementalMaterial.pdf).
- Supplemental dataset (SupplementalDataFile.xlsx).

## Data and code availability

All data is included in the figures and supplemental files. The code used to generate the main text figures, to recreate the predictions, and generate new predictions is available via GitHub (https://github.com/KULL-Centre/_2025_Larsen_degron_transcriptionfactors).

## Author contributions

F.B.L., N.J., K.E.J., F.D.E. and V.V. performed the experiments. F.B.L., V.V., K.E.J., N.J., K.L.-L., and R.H.-P. analyzed the data. K.L.-L. and R.H.-P. conceived the study. F.B.L., and R.H.-P. wrote the paper. All authors contributed to editing the paper and approved the final version before submission.

