## Supplemental Material for "Comprehensive degron mapping in human transcription factors"

**Fig. S1** *Transcription factor sublibrary flow cytometry profiles.* p.S2
**Fig. S2** *Transcription factor library perturbations.* p.S3
**Fig. S3** *Degron score correlations between biological and sorting replicates – Even Lib.* p.S4
**Fig. S4** *Degron score correlations between biological and sorting replicates – Odd Lib.* p.S5
**Fig. S5** *Degron score correlations between biological and sorting replicates – CT Lib.* p.S6
**Fig. S6** *Individual residue effects on tile degron scores.* p.S7
**Fig. S7** *C2H2 ZnF motifs are visible in heatmaps with fixed cysteine positions.* p.S8
**Fig. S8** *C2H2 ZnF motifs are mainly found in buried and structured regions.* p.S9
**Fig. S9** *C2H2 ZnFs are destabilized upon mutation of the Zn coordinating residues.* p.S10
**Fig. S10** *Degron score correlations between common TF library and cytosolic library tiles.* p.S11
**Fig. S11** *Predicted abundance partially explains CRX mutational effects in IDRs.* p.S12
**Fig. S12** *Low throughput flow cytometry gating strategy.* p.S13
**Fig. S13** *Fluorescence activated cell sorting gating strategy.* p.S14
**References for supplemental material** p.S15

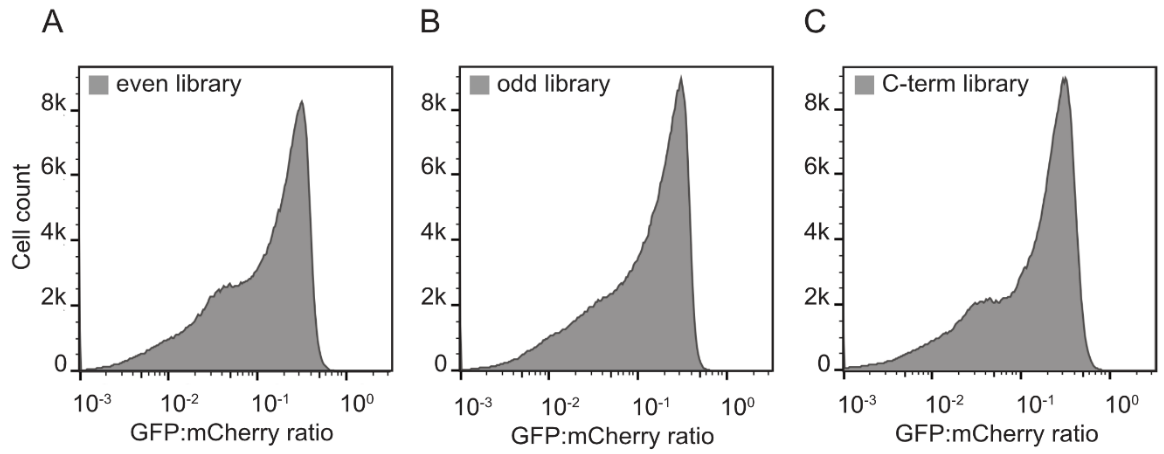


**Figure S1.** *Transcription factor sublibrary flow cytometry profiles*. Representative flow cytometry profiles displaying the quantification of the GFP:mCherry ratio of cells expressing the three human transcription factor sublibraries (even: n=4.71×$10^5$, odd: n=4.84×$10^5$ and C-terminal: n=4.76×$10^5$).

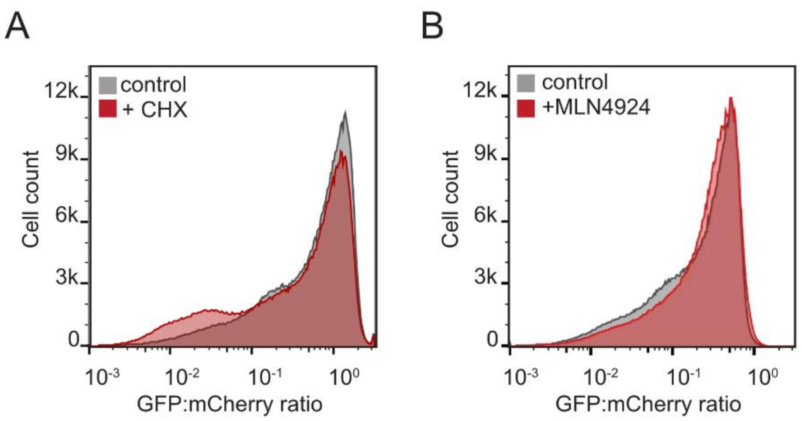


**Figure S2.** *Transcription factor library perturbations.* Representative flow cytometry profiles displaying the quantification of the GFP:mCherry ratio of cells expressing the TF odd sublibrary untreated (control, grey) (n=6×$10^5$), treated with 10 μg/mL cycloheximide (CHX) for 8 hours (n=6×$10^5$) to inhibit translation or 2 μM MLN4924 for 16 hours (n=6×$10^5$) to inhibit the NEDD8 E1 enzyme.

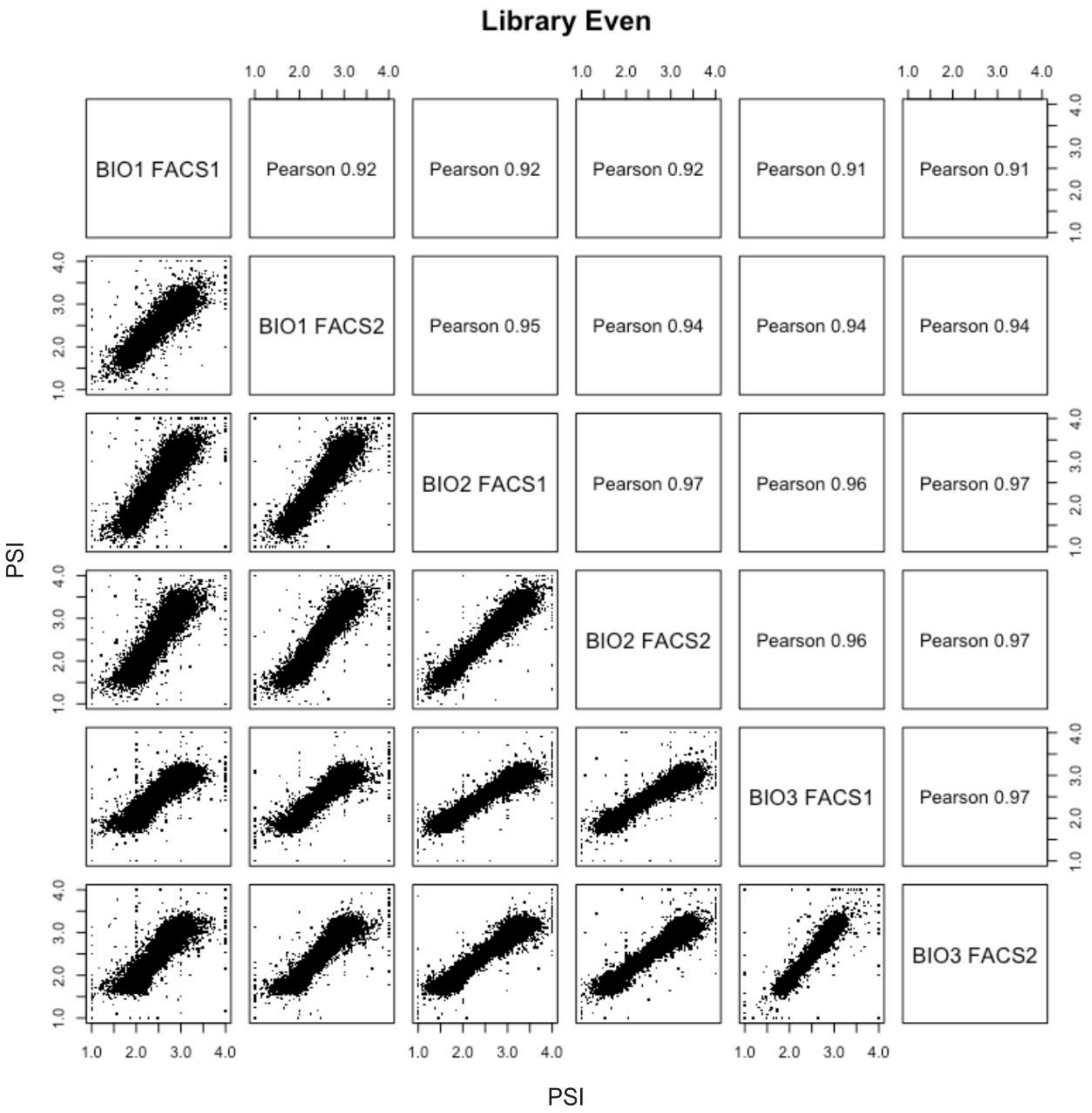


**Fig. S3** *Degron score correlations between biological and sorting replicates of the even TF library.* Individual correlations between the protein stability index (PSI) of all biological and sorting replicates. The Pearson correlation coefficient is indicated for each correlation plot (see methods).

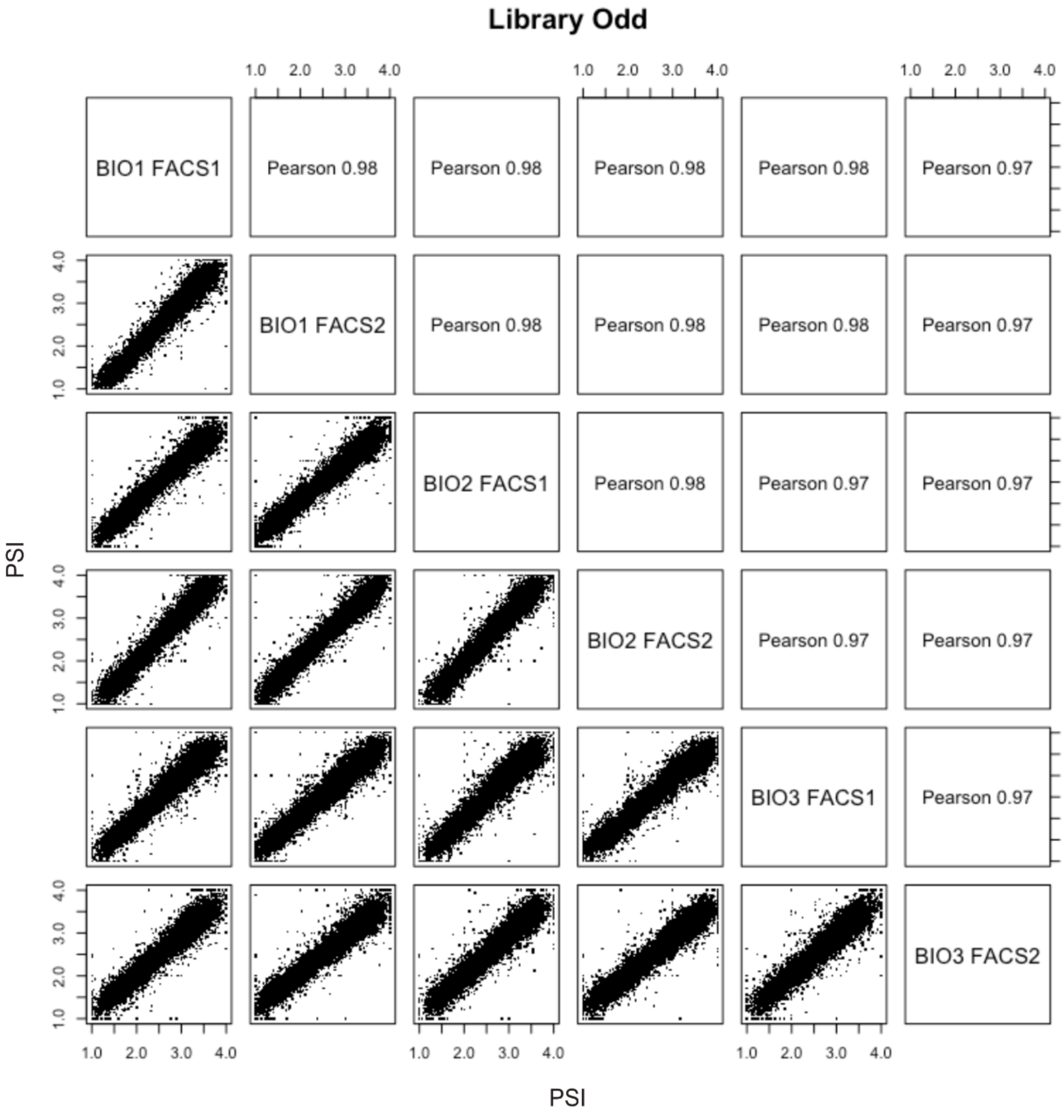


**Fig. S4** *Degron score correlations between biological and sorting replicates of the odd TF library.* Individual correlations between the protein stability index (PSI) of all biological and sorting replicates. The Pearson correlation coefficient is indicated for each correlation plot (see methods).

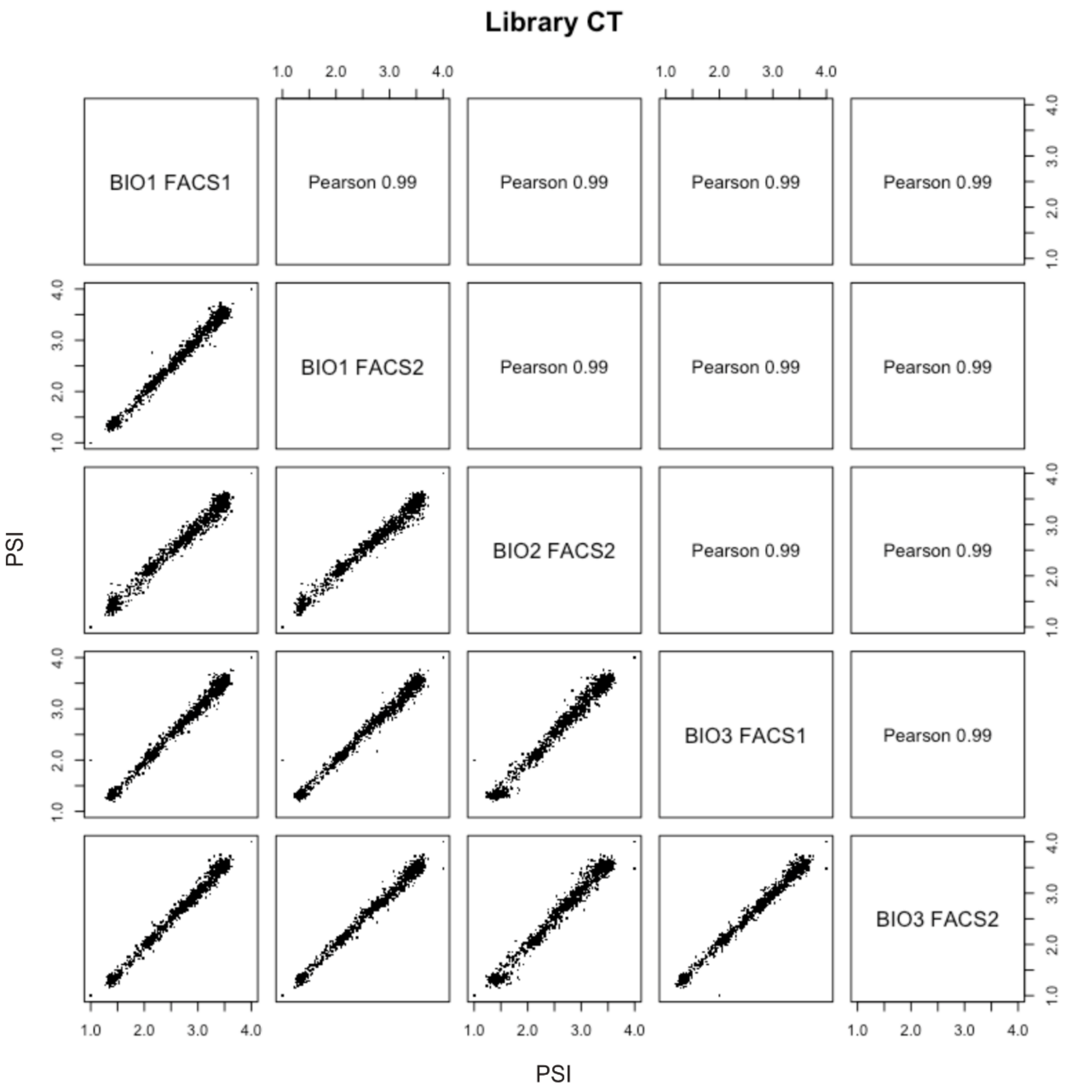


**Fig. S5** *Degron score correlations between biological and sorting replicates of the C-terminal (CT) TF library*. Individual correlations between the protein stability index (PSI) of all biological and sorting replicates. The Pearson correlation coefficient is indicated for each correlation plot (see methods).

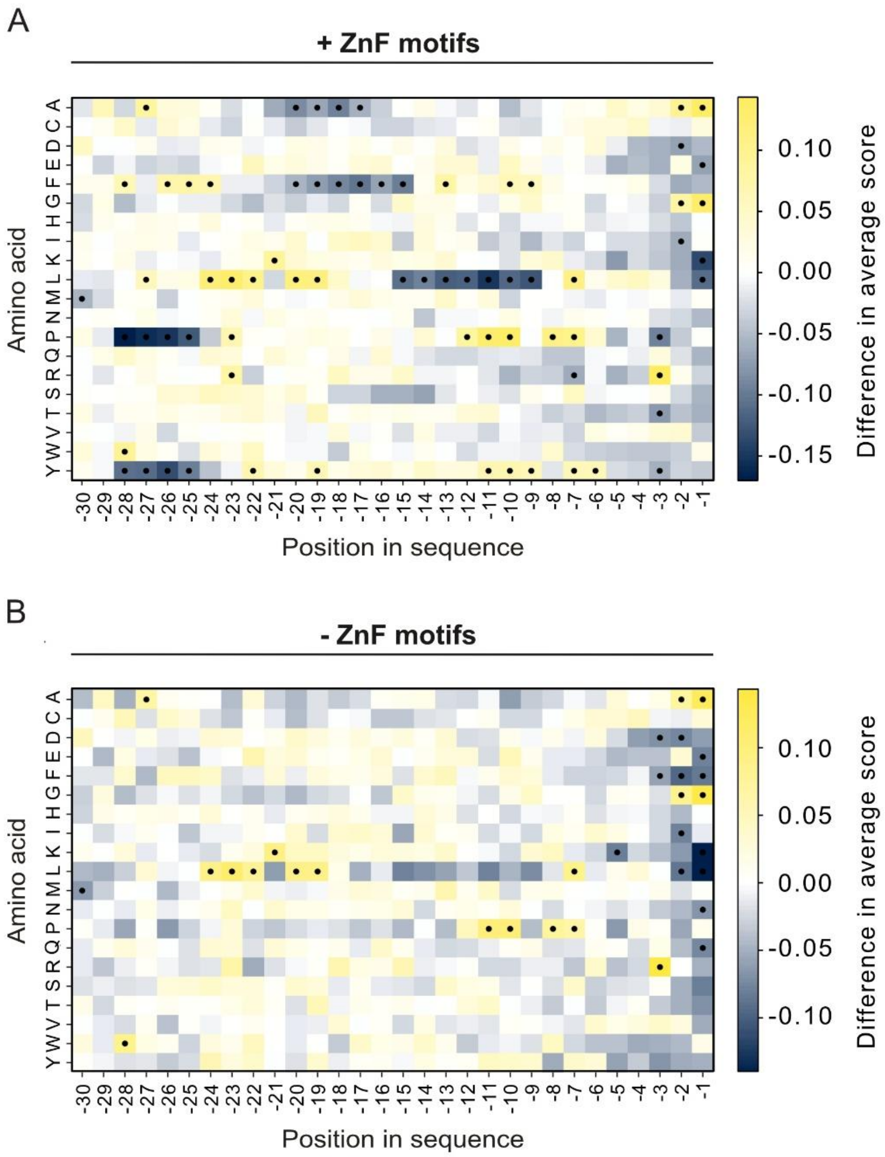


**Figure S6.** *Individual residue effects on tile degron scores.* Heatmaps displaying the difference in the average degron score of a single amino acid at a specific position from the average score of a single amino acid at any other position in the tile for all data either including (A) or excluding (B) tiles including an intact C2H2 ZnF motif. Yellow indicates an increase in the degron score, and dark blue indicates a decrease in the degron score. Black dots indicate statistical significance after Bonferroni correction based on Mann-Whitney U test ($p < 0.05/600$).

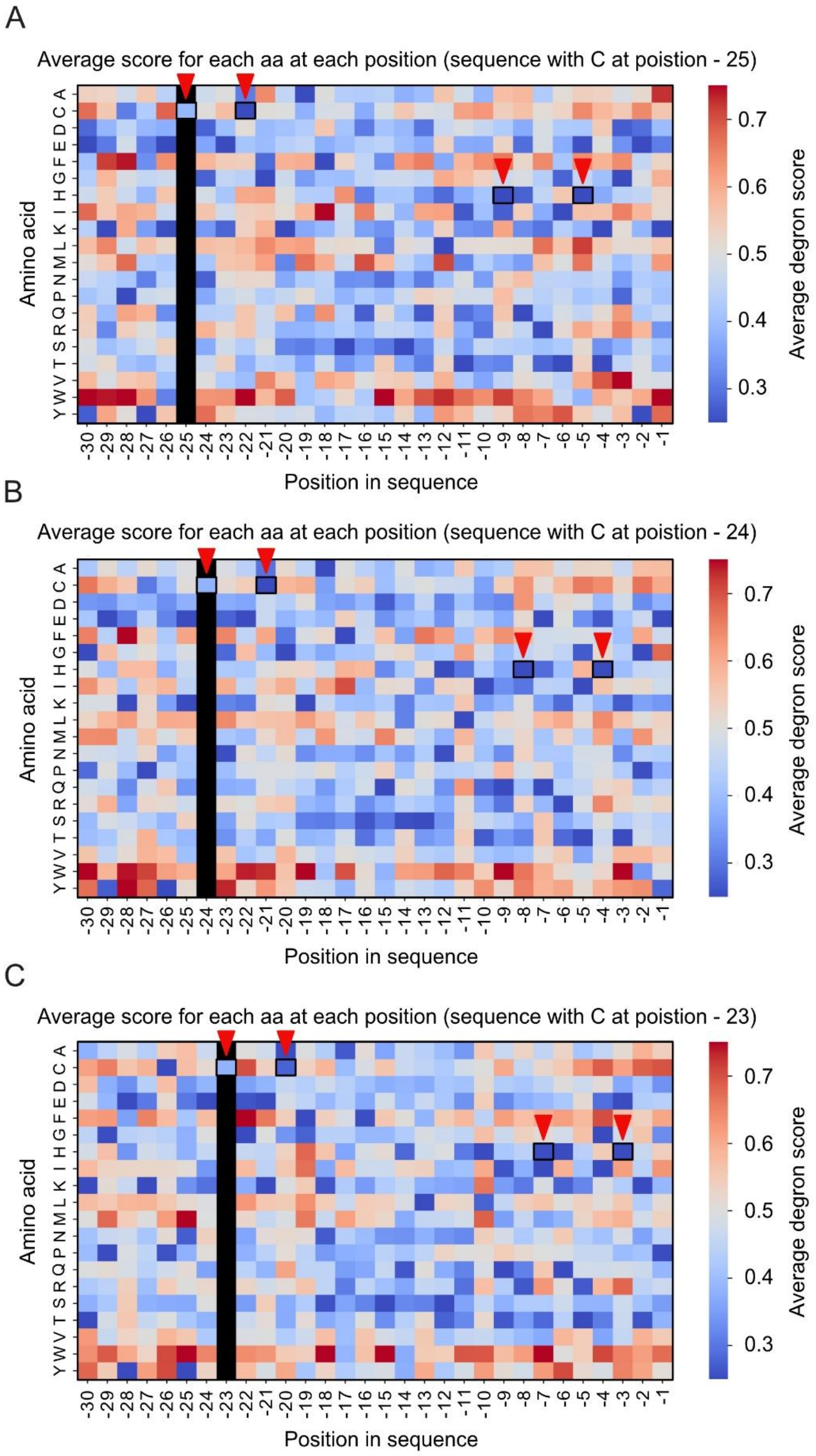


**Figure S7.** *C2H2 ZnF motifs are visible in heatmaps with fixed cysteine positions*. Heatmaps displaying the calculated degron score of a tile with a specific amino acid at a defined position when a cysteine is fixed at position (A) - 25, (B) - 24 or (C) - 23. Red arrows indicate the cysteine and histidine residues of the C2H2 ZnF motifs.

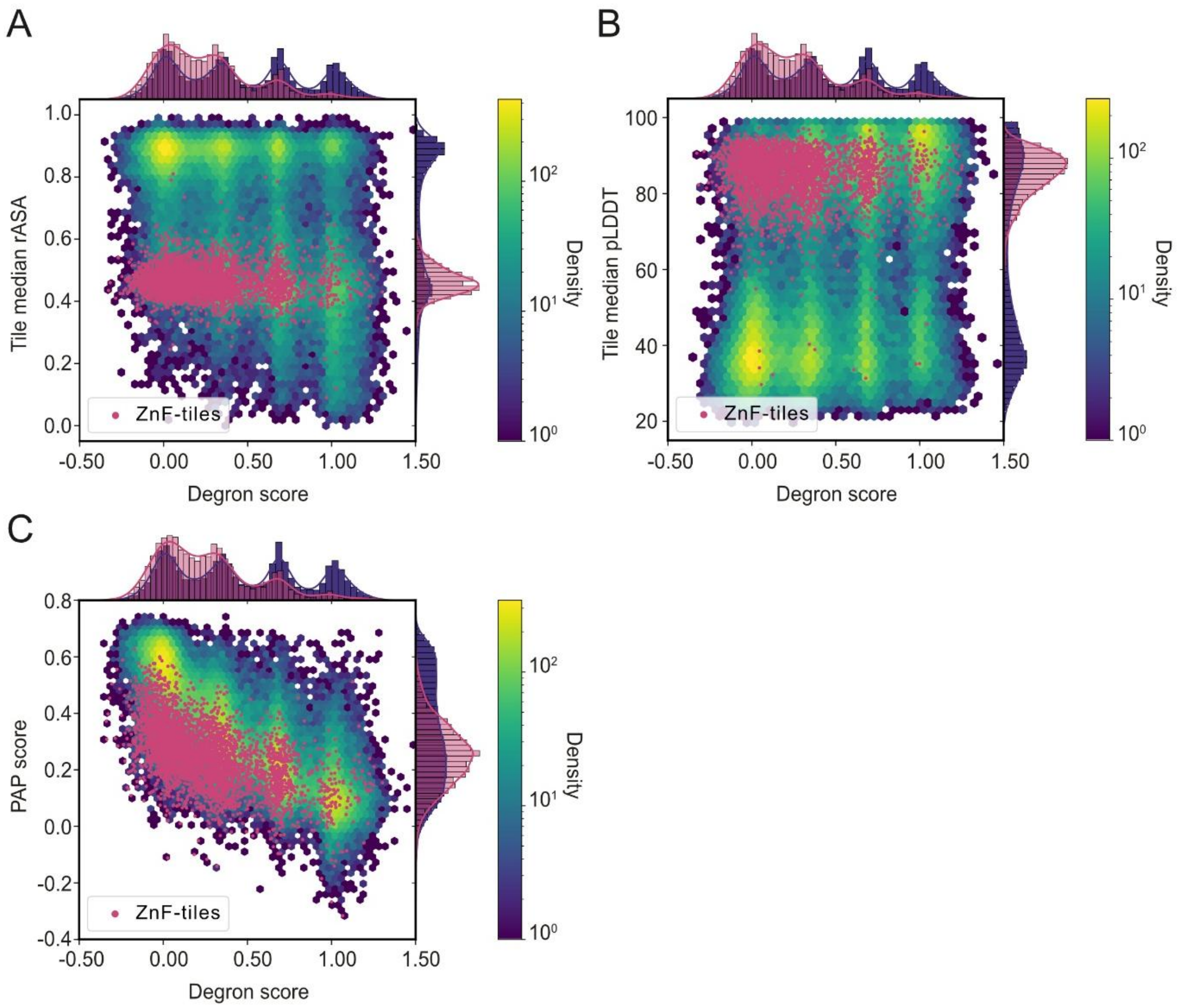


**Figure S8.** *C2H2 ZnF motifs are mainly found in buried and structured regions.* Correlation plots of human transcription factor library degron scores and per-tile median (A) relative accessible surface area (rASA) (absolute Pearson |r|: 0.41, absolute Spearman $|\rho|$: 0.39), (B) predicted local difference test (pLDDT) score (absolute Pearson |r|: 0.36, absolute Spearman $|\rho|$: 0.36), and (C) peptide abundance predictor (PAP) score (absolute Pearson |r|: 0.79, absolute Spearman $|\rho|$: 0.80), with tiles including an intact C2H2 ZnF motif marked in red. Yellow indicates high density, and dark blue indicates low density.

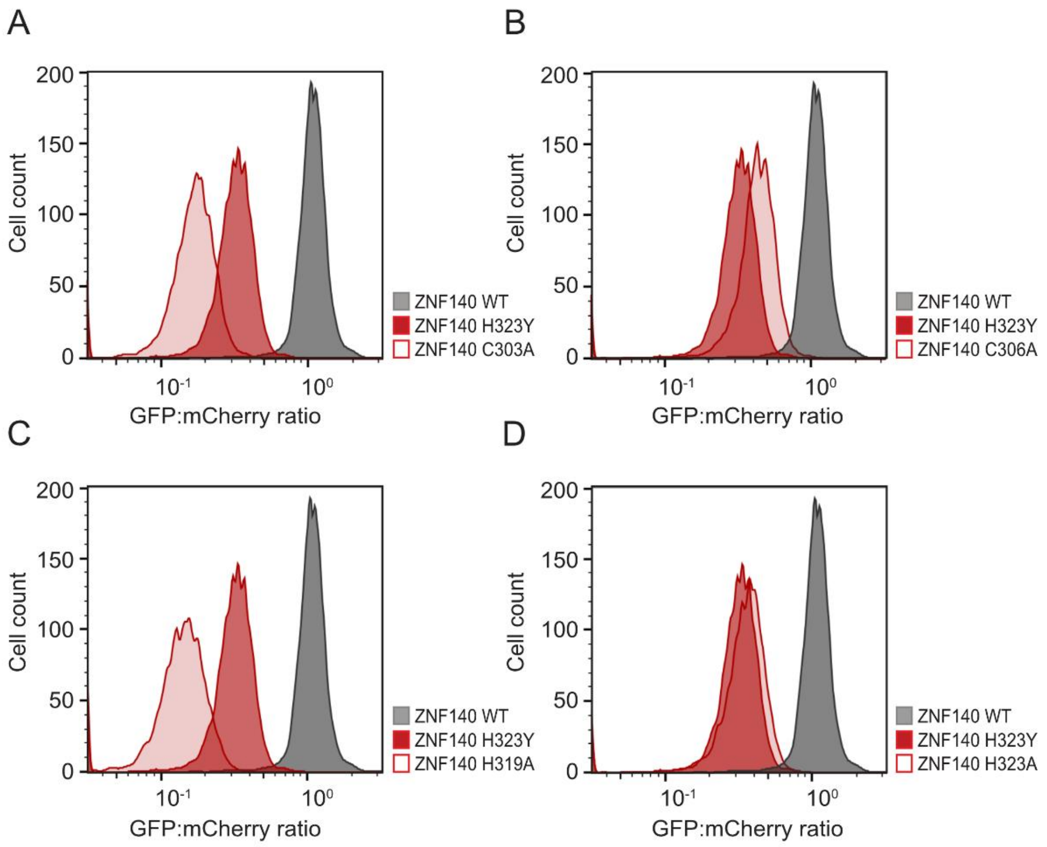


**Figure S9.** *C2H2 ZnFs are destabilized upon mutation of the Zn coordinating residues.* Representative flow cytometry displaying the quantification of the GFP:mCherry ratios of cells expressing tile 21 of ZNF140, including an intact ZnF motif as wild-type (WT) (n=5,089) and H323Y (n=5,210) for comparison, (A) C303A (n=5,265), (B) C306A (n=5,218), (C) H319A (n=5,120), and (D) H323A (n=5,070).

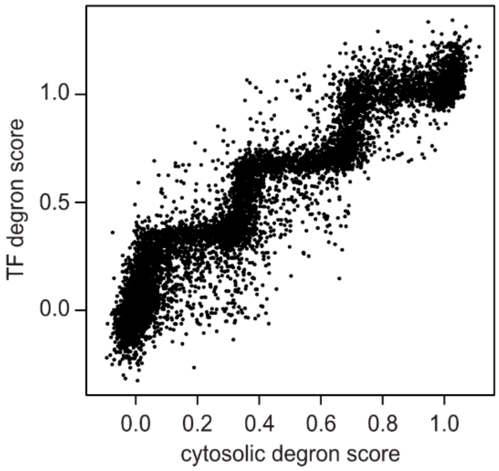


**Figure S10.** *Degron score correlations of common tiles from TF library and cytosolic library.* Correlation plot with degron scores of common tiles shared between the human transcription factor library and the human cytosolic library (8,645 tiles) (absolute Pearson |r|: 0.95). The step-like appearance of the plot is likely related to a misalignment of the four peaks in the two libraries due to the TF library containing fewer hydrophobic tiles (since the TFs do not contain large, folded protein domains). Fig. 5A shows the same data in a re-normalized version referred to as abundance score (see methods).

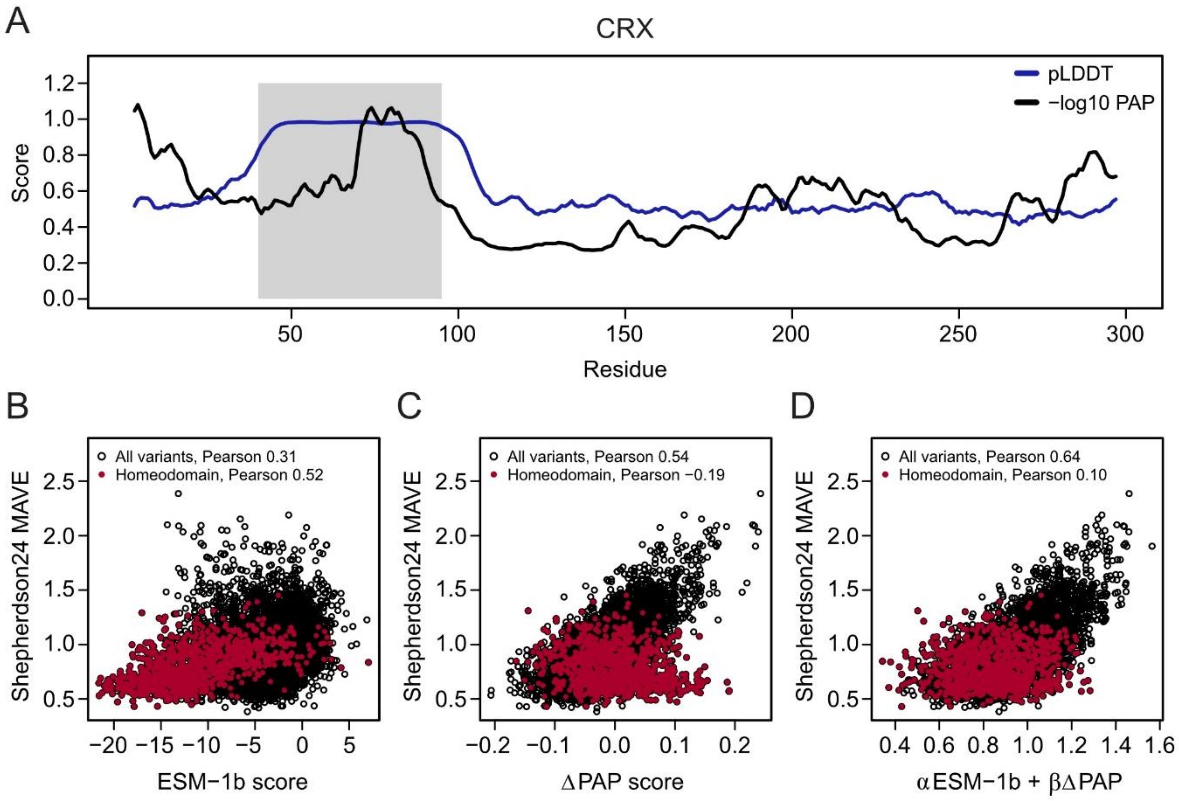


**Figure S11.** *Predicted abundance partially explains CRX mutational effects in IDRs.* (A) Profile of CRX (UniProt: O42186) displaying residue number on the x-axis and the -$\log_{10}$ peptide abundance prediction (PAP) score (black) and predicted local difference test (pLDDT) score derived from Alphafold2 (blue) on the y-axis (run. avg. size 5). The structured homeodomain (residues 40-95) is shaded. (B-D) Correlation plots displaying activity scores of CRX variants determined by Shepherdson *et al*. 2024 (1) on the y-axis and on the x-axis (B) variant effect scores calculated by the protein language model ESM-1b (2), (C) predicted variant peptide abundance score (ΔPAP) (3), and (D) optimized linear combination of ESM-1b and ΔPAP scores (α=0.02, β=2.65 and offset 1.06). Variants in the homeodomain are marked with red dots.

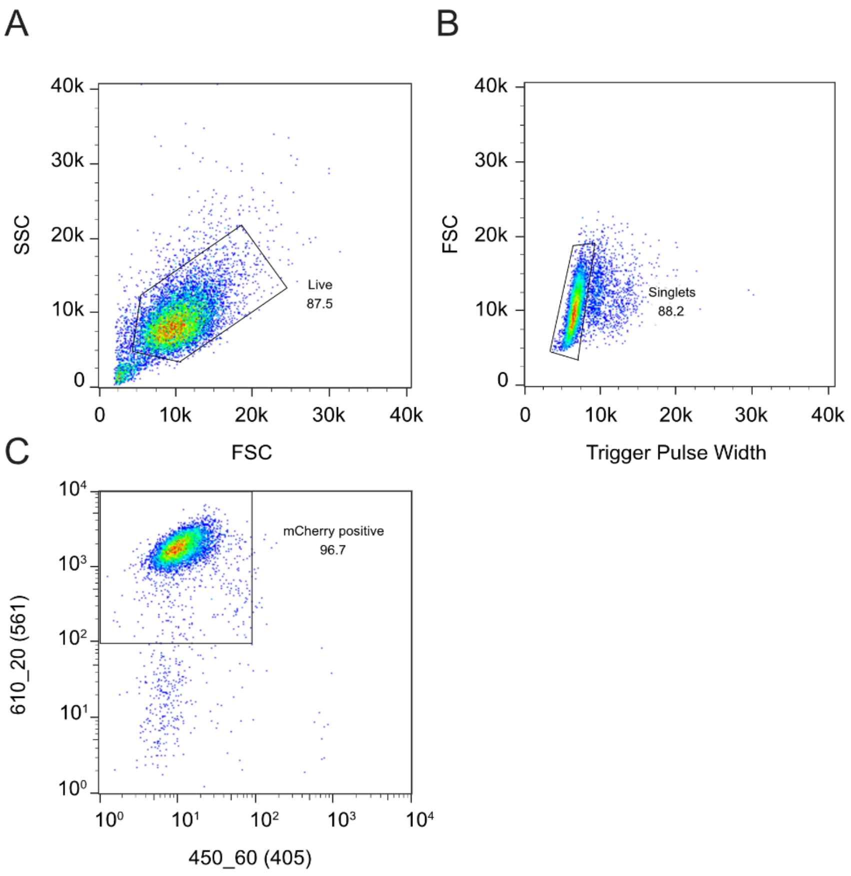


**Figure S12.** *Low throughput flow cytometry gating strategy.* The presented low throughput flow cytometry data were obtained by gating the population corresponding to (A) live cells based on the forward (FSC) and side (SSC) scatter. (B) Subsequently, single cells were gated from duplets based on the trigger pulse width and the FSC. (C) Finally, successfully recombined cells were isolated by gating cells that were BFP negative (450_60 (405)) and mCherry positive (610_20(561)).

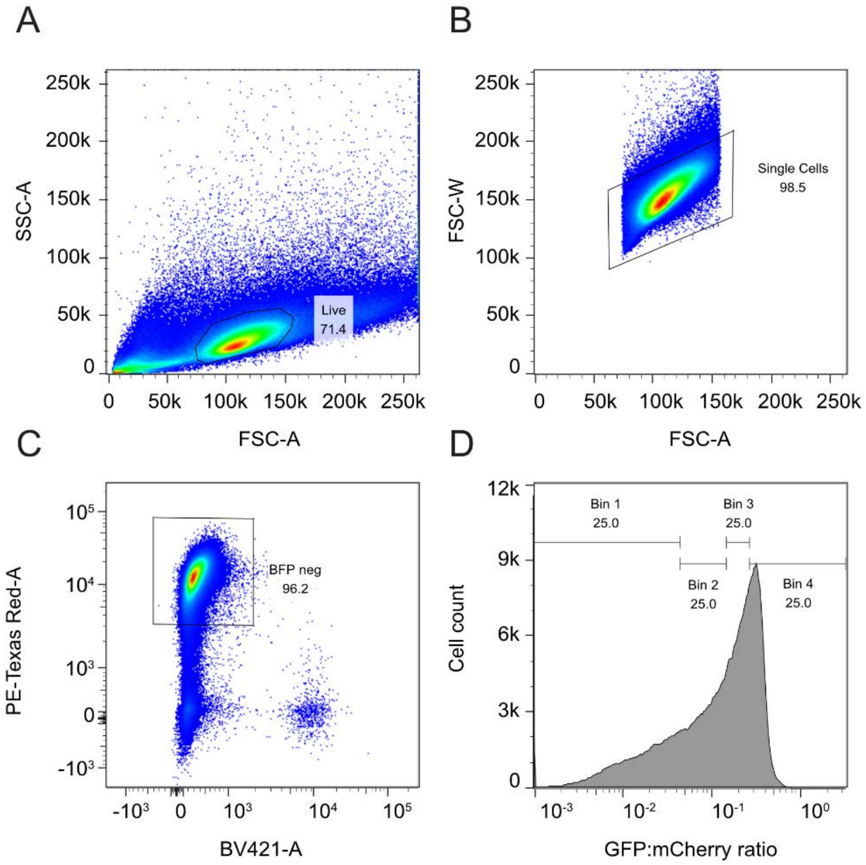


**Figure S13.** *Fluorescence activated cell sorting gating strategy*. Cells expressing the human transcription factor library were analysed and sorted by gating the population corresponding to (A) live cells based on the forward (FSC-A) and side (SSC-A) scatter. (B) Subsequently, single cells were gated from duplets based on the FSC-A and the FSC-W. (C) Successfully recombined cells were isolated by gating cells that were BFP negative (BV421-A) and mCherry positive (PE-Texas Red-A). (D) Finally, cells were sorted based on four gates (Bin 1-4), each containing 25 % of the cell population.
